# A bovine pulmosphere model and multiomics analyses identify a signature of early host response to *Mycobacterium tuberculosis* infection

**DOI:** 10.1101/2023.12.07.570553

**Authors:** Vinay Bhaskar, Rishi Kumar, Manas Ranjan Praharaj, Sripratyusha Gandham, Hemanta Kumar Maity, Uttam Sarkar, Bappaditya Dey

## Abstract

Interactions between the tubercle bacilli and lung cells during the early stages of tuberculosis (TB) are crucial for disease outcomes. Conventional 2D cell culture inadequately replicates the multicellular complexity of lungs. We introduce a 3D pulmosphere model for *Mycobacterium tuberculosis* infection in bovine systems, demonstrating through comprehensive transcriptome and proteome analyses that these 3D structures closely replicate the diverse cell populations and abundant extracellular matrix proteins, emphasizing their similarity to the *in vivo* pulmonary environment. While both avirulent BCG and virulent *M. tuberculosis*-infected pulmospheres exhibit commonalities in the upregulation of several host signaling pathways, distinct features such as upregulation of ECM receptors, neutrophil chemotaxis, interferon signaling, and RIG-1 signaling pathways characterize the unique early response to virulent *M. tuberculosis*. Moreover, a signature of seven genes/proteins, including IRF1, CCL5, CXCL8, CXCL10, ICAM1, COL17A1, and CFB, emerges as indicative of the early host response to *M. tuberculosis* infection. Overall, this study presents a superior *ex vivo* multicellular bovine pulmosphere TB model, with implications for discovering disease biomarkers, enabling high-throughput drug screening, and improving TB control strategies.

## Introduction

Tuberculosis (TB) in bovines poses a substantial challenge to global livestock economies and public health systems. *Mycobacterium bovis* and *M. tuberculosis* are the leading members of the Mycobacterium tuberculosis complex (MTC) causing bovine TB (BTB) [1, 2]. The recent addition of *M. orygis* to this group further emphasizes the critical risk of zoonotic transmission to humans [3, 4]. The presence of drug-resistant tubercle bacteria in both humans and cattle complicates TB eradication efforts, as cattle may act as reservoirs for drug-resistant strains [2, 5]. This underscores the need to address bovine TB as a pressing global public health priority [6].

In countries like India, where bovine TB incidence rates range from 2 to 50% among the cattle population, combating BTB is hindered by its complex epidemiology, challenges in implementing effective eradication policies, and the absence of suitable diagnostics and vaccines [7]. Additionally, the lack of species-specific disease models, and limited research infrastructure such as large animal biosafety level 3 (ABSL3) facilities, further constrain efforts to study and combat BTB. Establishing alternative disease models that replicate lung cellular environments and granulomatous TB pathology is crucial, particularly in settings with limited bio-containment resources, to broaden the scope of research on BTB pathogenesis. Current understanding of BTB pathogenesis heavily relies on live animal models and a limited range of *in vitro* models, primarily utilizing bovine peripheral blood mononuclear cells (PBMCs) and bovine macrophage cell lines [8–11]. In addition, mouse and human macrophage cell lines have also been employed to study host responses to bovine tubercle bacilli [12]. However, the variation in host responses among different species and mycobacterial strains underscores the importance of developing species-specific disease models to unravel intricate events during TB pathogenesis [13–16].

In recent years, three-dimensional (3D) lung culture systems have gained traction in studying TB and respiratory diseases due to their advantages over conventional two-dimensional (2D) lung cell culture methods [17–20]. These 3D cultures aim to mimic the anatomical, physiological, and functional features of the lung *in vitro*. While some 3D lung culture systems allow for the formation of cellular structures that resemble the architecture of lung tissue, including the presence of alveoli and airway epithelium, others offer improved multi-cellular cell-cell and cell-matrix interactions, allowing for enhanced cell differentiation, migration, and signaling [21–23]. These advancements enable researchers to investigate the interactions between lung cells and pathogens, environmental stimuli, and therapeutic compounds, leading to more accurate predictions and translation to pulmonary disease biology [24, 25].

In this study, we developed a 3D pulmosphere model using bovine primary lung cells to advance our understanding of BTB pathogenesis and host responses during early-stage infection [26]. By closely mimicking the lung microenvironment in terms of multicellularity, cell-cell, and cell-matrix interactions, this model offers a unique platform to study the intricate interactions between the tubercle bacilli and lung cells of the bovine host. Further, through comprehensive transcriptional and proteomic analyses, we identified critical signaling and metabolic pathways, genes, and proteins involved in the early stages of infection. This paves the way for discovering BTB disease biomarkers, correlates of TB immunity, and potential targets for host-directed therapies. In addition to providing a novel bovine 3D pulmosphere TB model, this study enhances our understanding of TB pathogenesis, contributing to the global goal of TB eradication through the discovery of improved prevention and control strategies.

## Results

### Development of bovine primary lung cell-derived 3D pulmospheres

To establish a 3D pulmosphere model for the study of *ex vivo* TB infection, first, the method for generation of pulmospheres from bovine primary lung mixed cells population was optimized followed by a thorough physiological characterization of the pulmospheres. **Fig 1A** depicts the overall workflow used for the generation of the 3D pulmosphere. To optimize the formation of uniform spheroids, four different seeding densities of the mixed primary lung cells per well were evaluated, including 1×10^4^, 1.5×10^4^, 2×10^4^, and 2.5×10^4^ cells in a U bottom 96-well TC plate. Notably, round-shaped, self-assembled, and reproducible spheroids were generated within 24 hours following centrifugation-based spheroid assembly (**Fig 1B**). The optimum seeding densities were found to be 1×10^4^ and 1.5×10^4^ cells per well, while a higher number of cells resulted in an improper assembly leading to irregularly shaped cell clumps. **Fig 1C** depicts the relative number of spheres generated at different cell seeding concentrations. These findings indicated that Poly-HEMA-coated U-bottom plates provided an appropriate microenvironment for pulmosphere development at the calibrated cell densities. Over a monitoring period of 28 days, the growth dynamics of bovine lung multi-cell spheroids were characterized. The mean diameter of the spheroids increased from 494 ± 70.58 μm at Day 7 post-sphere formation to 1205.4 ± 106.76 μm at Day 28 (n=20), revealing a substantial expansion in size over four weeks (**Fig 1D, E**). Further, the influence of primary lung cell culture passage number on pulmosphere formation was explored. We noted that the diversity and the heterogeneity of cell types underwent significant contraction following the second and the subsequent passages. Moreover, cells from passages zero and one lead to proper sphere formation, while cells from the latter passages reduced the success rate of spheroid formation as well as resulted in irregular-shaped cell clumps (**Supplementary Fig S1 A, B**). This relationship suggests a significant impact of the passage number on spheroid morphology and structure, underscoring the importance of early passages for maintaining the characteristics of 3D pulmospheres derived from primary lung cells.

**Figure 1.**
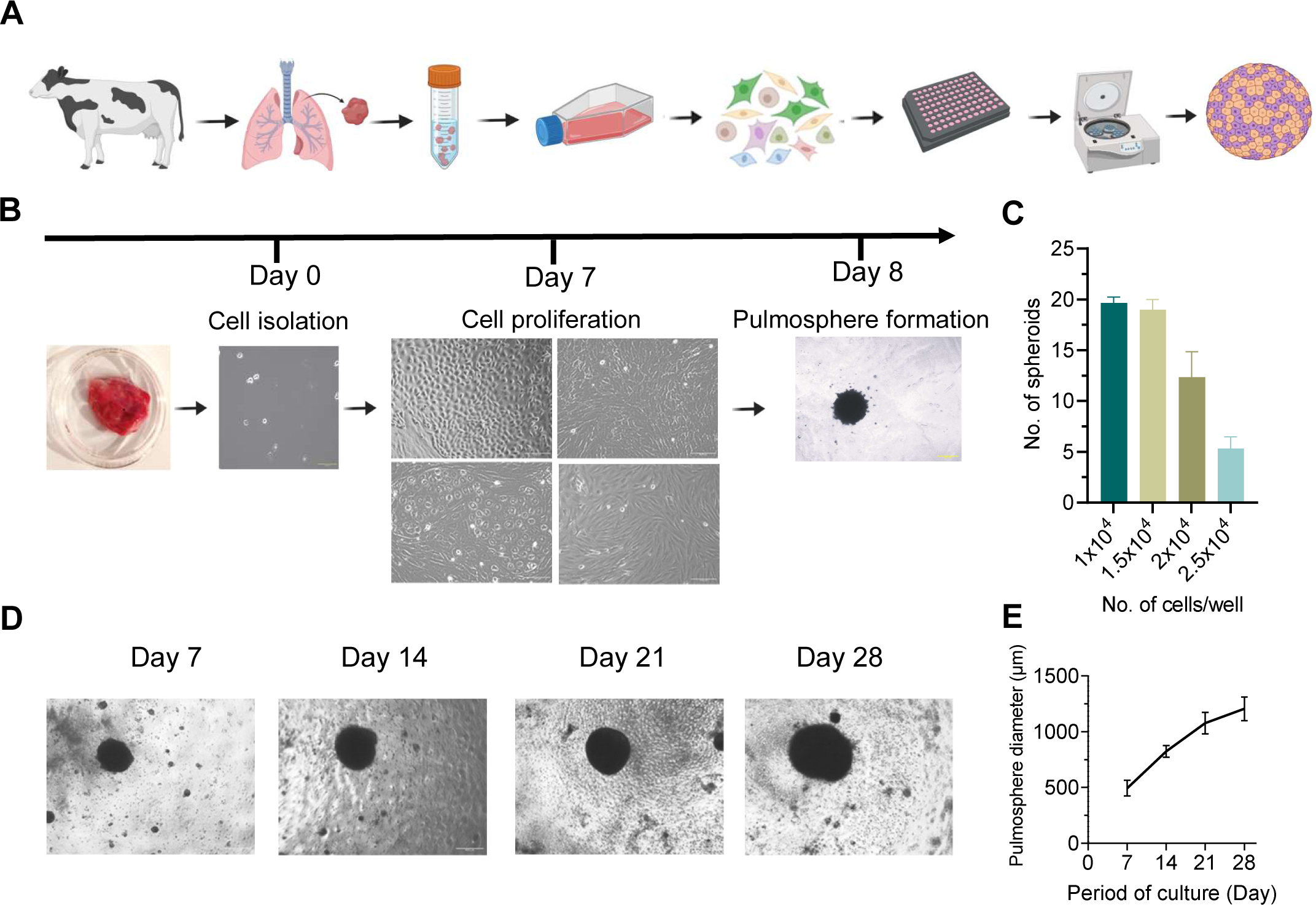
Generation and morphological characterization of bovine 3D pulmosphere. A Schematic cartoon representation of the protocol for bovine 3D pulmosphere development. B Bovine lung primary cells were isolated and grown for 7 days showing multiple cell types, pulmosphere was assembled and images were taken after 24 hours. Scale bars: 400 μm for 4x images (Day 0, day 7) and 200 μm for 10x images (Day 8). C Bar graph depicts the average number of compact pulmospheres formed with respect to the different cell seeding densities. D Representative images of pulmospheres captured at every week till 28 days. E The line diagram depicts the relative size of pulmospheres over the period of culture. Scale bars: 400 μm for 4x images. Data information: In (C, E) data are presented as mean ± sd. Statistical analysis included an unpaired two-tailed t-test for comparison between two. A 95% confidence interval or 0.05 threshold for significance (p < 0.05) was used in statistical tests

### 3D pulmospheres recapitulate pulmonary microenvironment over 2D monolayer culture

To investigate the cellular and functional features of the 3D pulmospheres we performed liquid chromatography-mass spectrometry (LC-MS) analysis of the whole protein extract 24 hours post-assembly of the pulmospheres. The proteomics workflow is provided in the **Supplementary Fig S2.** A pool of 2748 peptides was detected in the proteome of the pulmospheres (**Supplementary data file S1**), which were then subjected to global cell-type-specific enrichment analysis using the web-based tool WebCSEA [27], that unveiled the presence of 17 cell types (**Fig 2A)**. Further analysis of lung tissue-specific cell typing reveals 32 different types of lung cells encompassing epithelial, endothelial, fibroblast, pneumocytes, and an array of immune cells including macrophage, dendritic cells, neutrophil, T-cell, B-cells, plasma cells, etc. highlighting the complex multi-cellularity of the pulmonary environment (**Fig 2B**). A comparative assessment of protein expression patterns was performed between 2D monolayer cells, and 3D pulmospheres at 24 hours post-culture to understand the differences in cellular anatomy, physiology, and functional changes in the 3D pulmosphere compared to the 2D monolayer culture. **Supplementary Fig S3 A-C**, depicts the comparative omics analysis features: (A) Volcano plot, (B) PCA analysis, and (C) hierarchical clustering of peptide abundance) of the DEPs of 3D vs. 2D culture proteome.

**Figure 2.**
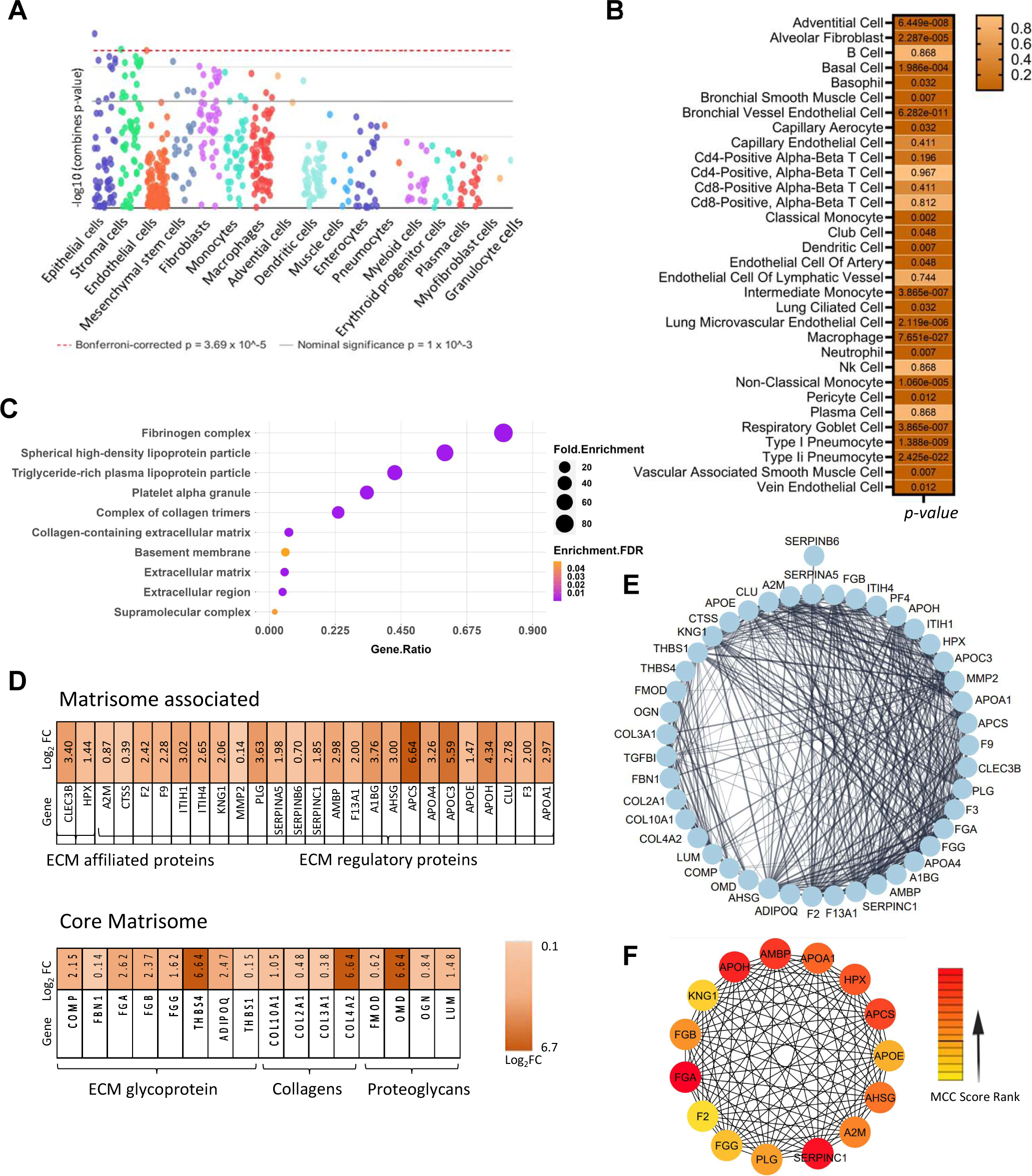
Cellular composition of 3D pulmosphere and key upregulated PPI networks over 2D culture. A Array of different cell types identified in bovine 3D Pulmosphere using WebCSEA. B Lung-tissue specific cell type enrichment analysis using ‘Tabula Sapiens’ single cell analysis showed the presence of 22 cell types as significantly enriched, p < 0.05. C Gene Ontology of cellular composition in 3D pulmosphere vs monolayer 2D lung cell culture. D Protein expression profile of ECM and associated proteins in 3D pulmosphere. E Densely interconnected protein network of ECM proteins derived via STRING analysis. F Top 15 critical hub genes involved in the structural organization of 3D pulmosphere were identified using Cyto-Hubba, a Cytoscape plug-in.

Strikingly, upon gene-ontology analysis (GO) of cellular components, a pronounced upregulation of the fibrinogen complex and collagen-containing extracellular matrix (ECM) proteins were found in the case of 3D pulmospheres (**Fig 2C**). Subsequent analysis of these proteins using a matrisome database revealed a strong association of the majority of the core matrisome and matrisome-associated proteins in the case of 3D pulmospheres [28] (**Fig 2D**). This finding suggests a fundamental shift in the protein expression profile that emphasizes the significance of the 3D architecture in promoting the deposition and assembly of ECM components, which are integral to lung structure and function. Further insight into the upregulated ECM network within the 3D pulmosphere was gained through the identification of closely interconnected ECM proteins by STRING analysis (**Fig 2E**). The top 15 major ECM proteins showing a higher number of interconnected nodes were APOH, APOA1, HPX, AMBP, APCS, APOE, AHSG, A2M, SERPINC1, PLG, FGG, F2, FGA, FGB, KNG1 (**Fig 2F)**. In the context of lung ECM formation, FGA, FGB, and FGG contribute to the ECM assembly through fibrin formation and provide structural integrity of the ECM. A2M and KNG1 are associated with protease inhibition, potentially regulating ECM remodeling processes. APOA1, APOE, and HPX are involved in lipid metabolism and transport, which can influence ECM stability. AMBP, APCS, APOH, and AHSG contribute to ECM formation by interacting with ECM proteins and also contribute modulation of inflammation and cell adhesion response. This intricate web of ECM molecules highlights the dynamic interactions and organization that contribute to the structural integrity and functionality of the pulmosphere. In addition to the upregulation of ECM proteins, several biological pathways were upregulated in the 3D pulmosphere (**Supplementary data file S2**). These include tissue homeostasis, epithelial cell proliferation, positive regulation of cell development, regulation of vasculature development, response to wounding, humoral immune response, negative regulation of apoptotic signaling pathway, and response to oxidative stress, etc. indicate that our 3D model not only captures the diverse cell types present in the lungs but also represents the complex extracellular matrix milieu, and the cellular behavior and tissue homeostasis pivotal for multicellular organismic development. This collective presence of distinct cell populations and ECM proteins underscores the physiological relevance of the 3D model in recapitulating the complexity of the *in vivo* lung environment.

### Development of 3D pulmosphere TB disease model

The establishment of a bovine 3D pulmosphere TB disease model represents a significant milestone, providing a platform for comprehensive investigations into the intricate interactions between *M. tuberculosis* or *M. bovis* and bovine lung cells within a physiologically relevant context. To construct the bovine TB infection model, both virulent *M. tuberculosis* and the vaccine strain *M. bovis* Bacille Calmette-Guérin (BCG) were employed. The inclusion of the vaccine strain enabled a comparative analysis of the host response elicited by a virulent *M. tuberculosis* strain versus an avirulent *M. bovis* BCG strain. For real-time monitoring of infection dynamics, fluorescent protein-expressing variants of *M. tuberculosis* and *M. bovis* BCG were used [29]. **Fig 3A** depicts the workflow employed for the generation of the 3D pulmosphere TB infection model. Seven days following the culture of bovine primary lung cells, the cells were subjected to infection with a pre-calibrated multiplicity of infection (MOI) of 1:10. This time point was chosen to harness the maximal heterogeneity of cell types inherent to early-passage of primary lung cells, ensuring an optimal representation of the multicellular pulmonary microenvironment in the development of the disease models. After infection, lung cells were assembled into 3D pulmospheres using the same methodology established for the formation of uninfected 3D pulmospheres. The infected 3D pulmospheres were monitored over 21 days through live fluorescence microscopy at weekly intervals **(Fig 3B)**. This longitudinal observation unveiled distinctive infection patterns between the attenuated vaccine strain *M. bovis* BCG and the virulent *M. tuberculosis* strain. The BCG strain exhibited restrained replication within the pulmospheres throughout the three-week study, predominantly localizing to focal infection. In stark contrast, the virulent *M. tuberculosis* strain demonstrated unhindered replication throughout infection and displayed widespread dissemination to all regions of the pulmospheres from the initial foci of infection **(Fig 3B)**. The successful establishment of the bovine 3D pulmosphere TB infection model has ushered in the ability to visualize the progression of infection dynamics *ex vivo*. This dynamic and physiologically representative system provides a unique window into the complex interplay between mycobacterium species and bovine lung cells, facilitating further studies to understand the deeper insights into the mechanisms underpinning TB pathogenesis.

**Figure 3.**
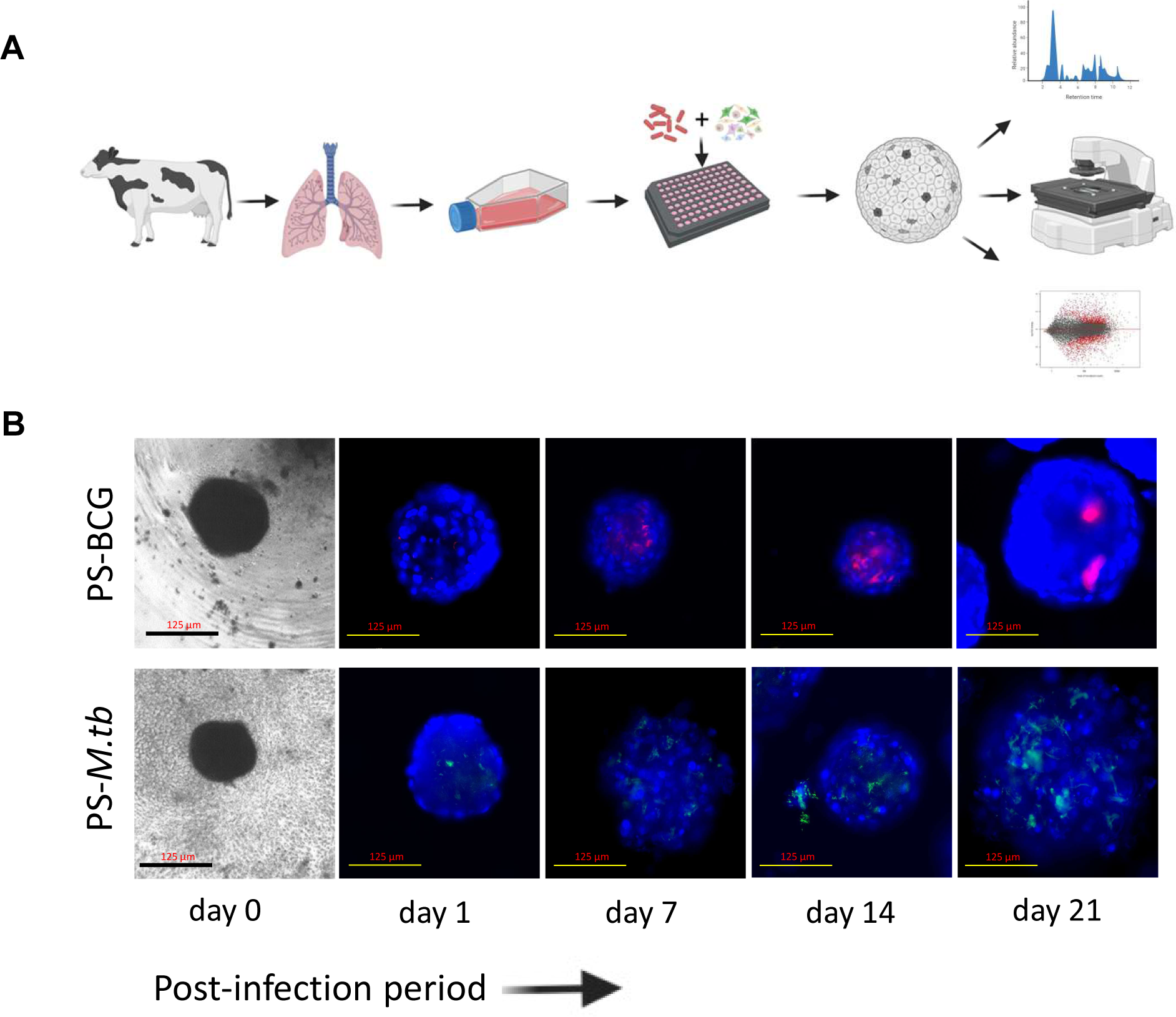
Establishment of bovine 3D pulmosphere TB infection model. A Schematic cartoon representation of the protocol for the establishment of bovine 3D pulmosphere infection and downstream analysis. B Seven-day cultures of bovine primary lung cells were infected with either *M. bovis* BCG-mCherry or *M. tb*-eGFP at an MOI of 1:10 and pulmospheres were assembled (day 0), and monitored under fluorescence microscope weekly till 21 days. The figures depict representative photographs of infected pulmospheres at different time points. Scale bars: 125 μm for 10x images.

### Transcriptomic and proteomic analysis of host response to BCG and *M. tuberculosis* infection in the bovine 3D pulmosphere model

To decipher the intricate host-pathogen interactions occurring at the early phase of infection within the bovine pulmonary environment, we analyzed the global transcriptomic and proteomic patterns of the infected and uninfected pulmospheres at 24 hours post-sphere formation. A stringent analytical framework was employed, wherein genes and proteins demonstrating at least a 2-fold change, and with an adjusted p-value of <0.05 were deemed differentially expressed (DE). The proteome and transcriptome data analysis workflow were provided in **Supplementary Fig S2 and S4,** respectively**. Supplementary Fig S5 A-D** and **S6 A-E** depict the comparative omics analysis features (Volcano plot, PCA analysis, and hierarchical clustering) of the transcriptome data for PS-BCG and PS-*M. tb* compared to uninfected pulmospheres, respectively. Comparative analysis revealed that the bovine 3D pulmospheres exhibited substantial transcriptional and proteomic responses upon infection with *M. bovis* BCG and *M. tuberculosis* strains **(Fig 4,5**). Specifically, 766 differentially expressed genes, and 281 differentially expressed proteins were identified in *M. bovis* BCG-infected pulmospheres, whereas 2129 DEGs and 402 DEPs were detected in *M. tuberculosis*-infected pulmospheres, relative to uninfected counterparts indicating that a greater number of genes exhibited altered expression levels in response to the virulent *M. tuberculosis* challenge, both at the transcript and protein levels (**Fig 4A, B**). Further, compared to BCG infection 3828 genes and 406 proteins were differentially expressed in the case *of M. tuberculosis* infection (**Fig 4A, B**). This finding underlines the intricate and multifaceted nature of the host response to *M. tuberculosis* infection when compared with the avirulent *M. bovis* BCG-induced response. Additionally, a comparison with the publicly available transcriptome data from a variety of *M. tuberculosis* infection models employed by several previous studies revealed that the number of DEGs in our 3D pulmosphere model was notably higher following *M. tuberculosis* infection (**Fig 4C**) [8, 30–34]. This finding highlights the 3D pulmosphere as a more robust system compared to the single-cell, or PBMC-based models representing an enormous depth of complexity and cellular responses induced during *M. tuberculosis* infection.

**Figure 4.**
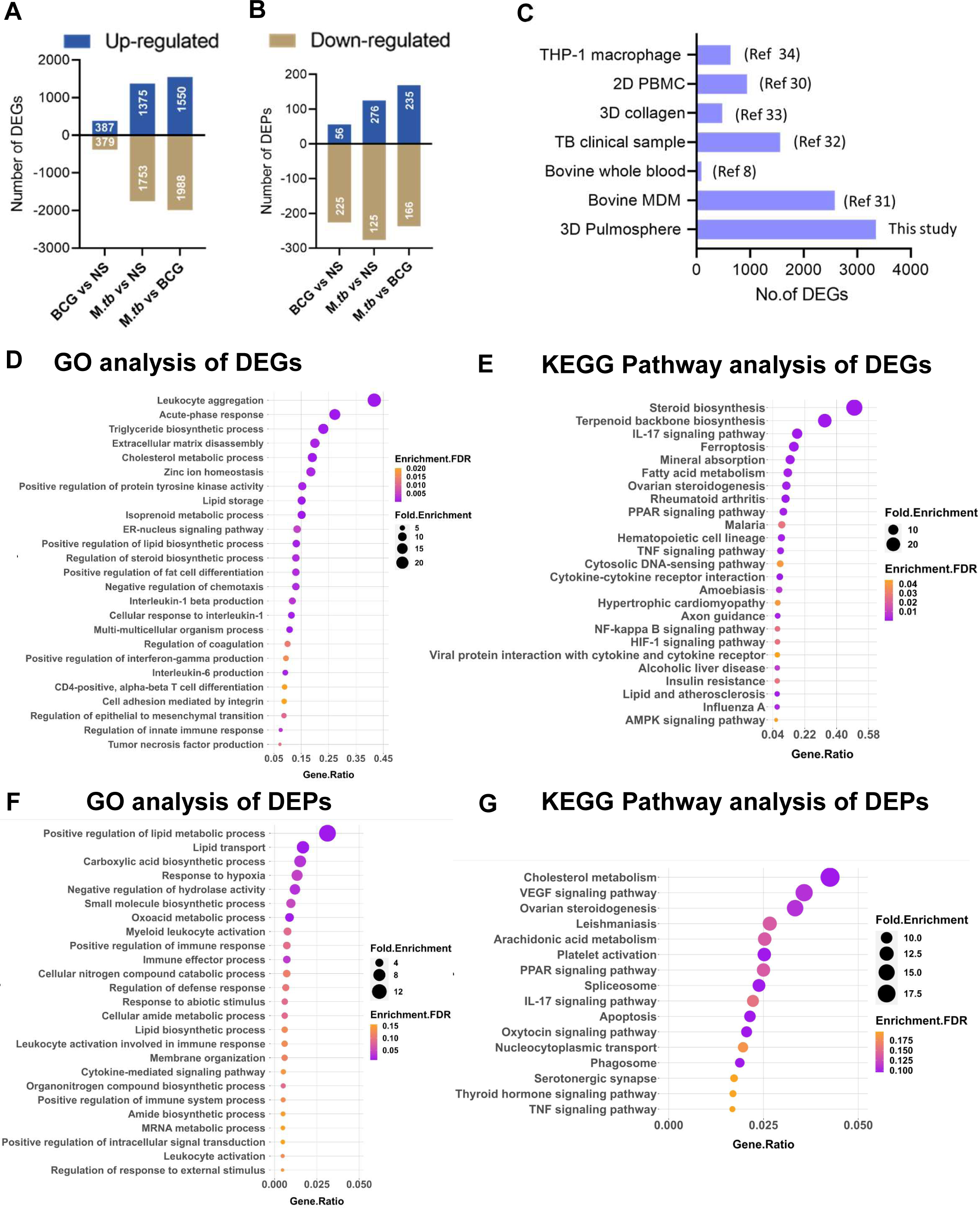
Differential gene, protein expression, and functional enrichment analysis of the DEGs and DEPs in 3D pulmospheres following BCG and *M. tuberculosis* infection. A No. of differentially expressed proteins in BCG and *M. tb* infected pulmospheres. B No. of differentially expressed genes in BCG and *M. tb* infected pulmospheres. C Comparative view of the DEG no. in PS-*M. tb* (this study) and selected TB infection model reported previously. The numbers in the parenthesis are the corresponding references. D, E Biological processes (D), and KEGG pathway analysis of DEGs (E) of PS-BCG over uninfected pulmospheres. FDR < 0.05, and gene number >5 in a pathway were used as cutoff values. F, G Biological processes (F), and KEGG pathway analysis of DEPs (G) of PS-BCG over uninfected pulmospheres, respectively. FDR < 0.05, and gene number >5 in a pathway were used as cutoff values.

**Figure 5.**
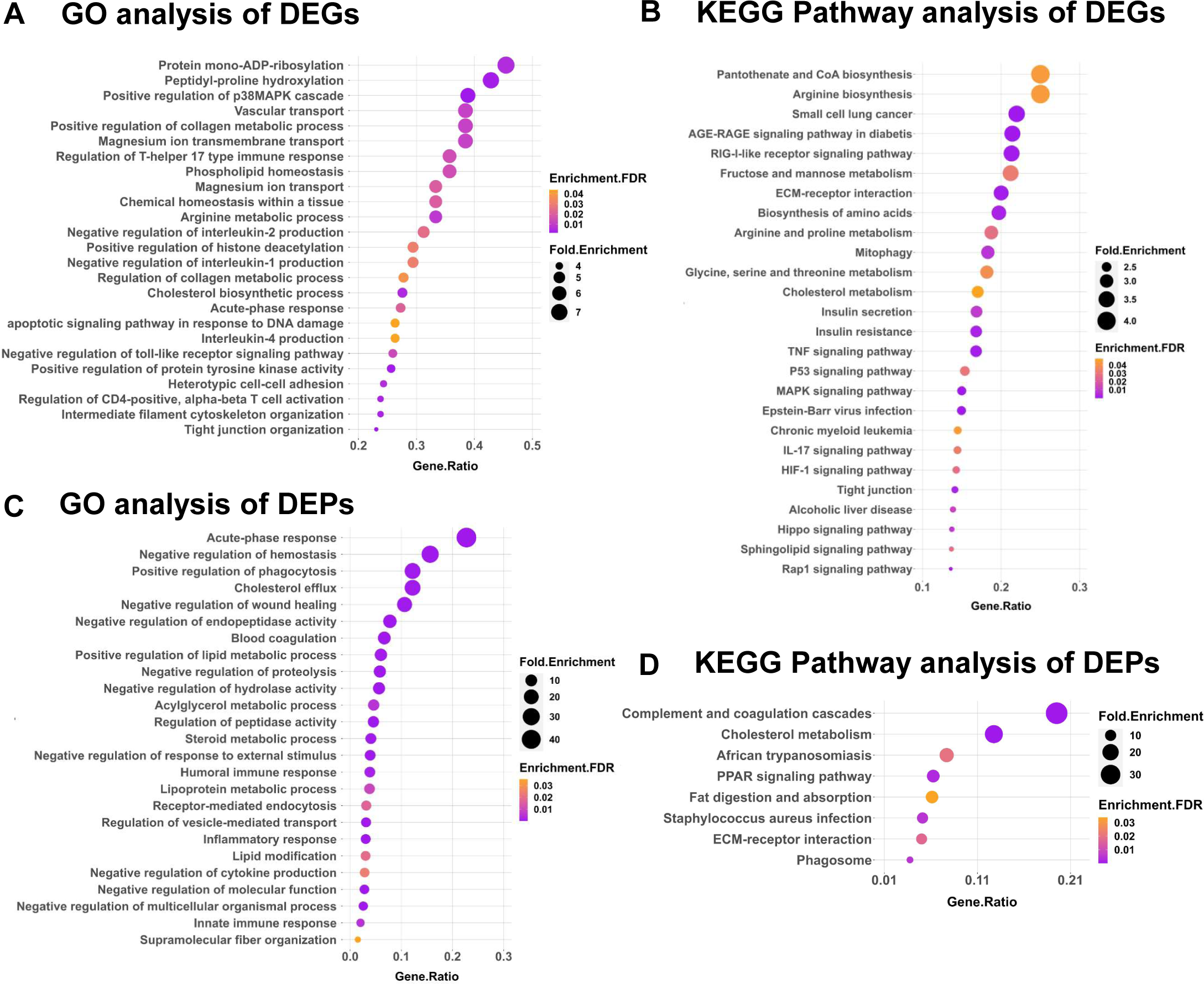
Functional enrichment analysis of the up-regulated genes and proteins in case of *M. tuberculosis* infection vs. uninfected pulmospheres. A, B Biological processes (A) and KEGG pathway (B) analysis of DEGs of PS-*M.tb* over uninfected pulmospheres FDR < 0.05, and gene number >5 in a pathway were used as cutoff values. C, D Biological processes (A) and KEGG pathway (B) analysis of DEPs of PS-*M.tb* over uninfected pulmospheres FDR < 0.05, and gene number >5 in a pathway were used as cutoff values.

### Functional enrichment analysis of differentially expressed genes following M. tuberculosis and M. bovis BCG infection

To gain insights into the biological implications of the observed differential expression, we performed GO and Kyoto Encyclopaedia of Genes and Genomes (KEGG) pathway enrichment analyses on upregulated and downregulated genes and proteins. Functional enrichment of the upregulated DEGs into the biological process via GO analysis reveals three major themes within the top-25 GO terms: immune response, cellular organization and ECM modification, and lipid metabolism were up-regulated in BCG-infected pulmosphere compared with the uninfected pulmosphere (**Fig 4D**). Pathways related to activation of innate immune responses, IFN-γ, TNF-α, IL-1β, IL-6 production, leukocyte aggregation, and acute phase response were upregulated in the case of BCG infection. In contrast, clusters of genes belonging to leukocyte activation, differentiation, proliferation, and chemotaxis were downregulated (**Supplementary Fig S7 A)**. Further, KEGG pathway analysis of the DEGs in the BCG group showed additional upregulation of IL-17, TGF-beta, Cytosolic DNA sensing pathway, PPAR, and NF-kappa B signaling pathways as major immune-related pathways (**Fig 4E**). In contrast, the top-down regulated immune pathways were complemented and coagulation cascades, neutrophil extracellular trap formation, Rap 1 signalling pathway, and Toll-like receptor signaling pathway (**Supplementary Fig S7 B**).

Functional enrichment via GO analysis of the DEPs in the BCG-infected pulmospheres reveals significant upregulation of similar immune response-related genes and lipid biosynthetic pathways as observed in the case of DEGs, however, several additional metabolic pathways such as carboxylic acid biosynthesis, amide metabolism, and mRNA metabolic processes, etc. were found to be upregulated (**Fig 4F**). However, KEGG pathway analysis didn’t show any significantly upregulated pathway (**Fig 4G).** GO analysis of the downregulated proteins showed oxidative phosphorylation, AGE-RAGE signaling, Th17 differentiation, and NOD-like receptor signaling pathway as the major enriched pathways (**Supplementary Fig S8 A**). KEGG analysis of DEPs identified downregulation of pathways related to several cancer types, phagocytosis, adipocytokine signaling, and RIG-I-like receptor signaling (**Supplementary Fig S8 B**).

On the other hand, in the case of *M. tuberculosis* infection, the immune response related to GO-BP terms shows upregulation of Th-17, Th-2 (IL-4), acute phase response, and genes related to negative regulation of IL-1, IL-2 production, and apoptotic signaling pathway (**Fig 5A**). Additionally, the KEGG pathway analysis indicated AGE-RAGE signaling pathway, RIG-1-like receptor signaling pathways, ECM-Receptor interaction, mitophagy, TNF and IL-17 signaling pathways, MAPK Signaling, RAP-1, and HIF-1 signaling pathways as some of the leading upregulated pathways in case of *M. tuberculosis*-infected pulmosphere (**Fig 5B**). Whereas, the downregulated pathways include complement and coagulation cascades, cell cycle process, ribosome biogenesis pathways, and neutrophil extracellular trap formation pathways (**Supplementary Fig S9 A**). KEGG analysis identifies mitochondrial respiratory chain complex assembly as one of the major down-regulated pathways (**Supplementary Fig S9 B**).

Functional enrichment via GO analysis of the DEPs in the case of *M. tuberculosis* infection led to the upregulation of several biological processes under GO-BP terms including acute phase response, Cholesterol, and lipid metabolism, positive regulation of phagocytosis, negative regulation of wound healing, humoral immune response, inflammatory and innate immune response (**Fig 5C**), However, KEGG pathway analysis reveals upregulation of additional pathways, such as the complement and coagulation cascades, PPAR signalling pathway, and ECM-Receptor interaction in case of *M. tuberculosis* infection (**Fig 5D**), Notably, both GO-BP terms and the KEGG pathways involving the downregulated genes remain similar to that of the DEGs, respectively (**Supplementary Fig S10 A, B**).

### Protein interaction network, hub genes, and functional clusters

Next, to have an in-depth understanding of the functional interactions among DEGs, we constructed protein-protein interactions (PPI) maps using STRING software, and identified the potential interacting network of proteins (**Supplementary data file S3**). The significantly interacting proteins from STRING analysis were subsequently analyzed via the molecular complex detection (MCODE) algorithm using Cytoscape to identify the key fundamental process modulated during infection. A total of ten clusters were identified among the upregulated genes of BCG-infected pulmospheres, whereas 36 clusters were identified among the genes of *M. tuberculosis*-infected pulmospheres. The top up-regulated cluster in the case of BCG infection was identified with an MCODE score of 19.8, whereas in the case of downregulated genes, 14 clusters were found with the highest score of 31.43 (**Supplementary data file S4**). To investigate the biological and immunological behavior of the highly interconnected upregulated modules, we performed GO enrichment using ClueGo on the top two MCODE clusters shown in **Fig 6A**, and **Fig 6B**. Subsequently, ClueGo analysis reveals sterol biosynthetic process, isoprenoid biosynthetic process, cellular response to interleukin-1; and neutrophil migration, regulation of macrophage activation and positive regulation of type-2 immune responses were the major networks in the upregulated top two highly interconnected gene clusters during BCG infection (**Fig 6C, D)**. Further, Cytohubba analysis reveals HMGCR, HMGCS1, SQLE, FDT1, FDPS, MSMO1, CYP51A1, DHCR24, MVD, and LSS are the key genes among all the upregulated genes in the case of BCG infected pulmospheres (**Fig 6E**).

**Figure 6.**
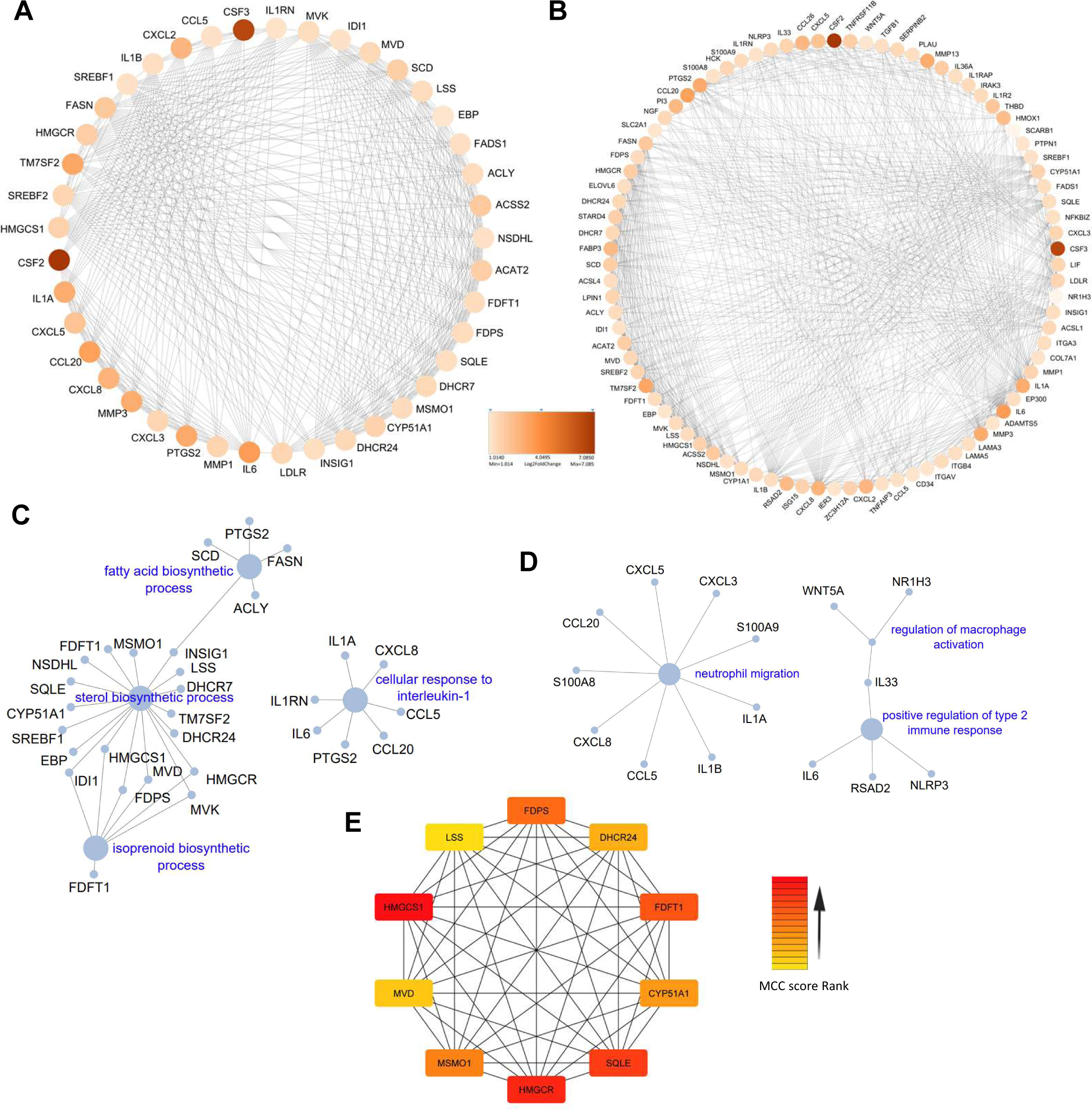
Network analysis of the up-regulated genes in BCG infected pulmospheres. A, B Critical networks cluster1 (A) and cluster2 (B) were constructed using MCODE algorithm in cytoscape. C, D ClueGO based GO of biological process, and immune related pathways in highly interconnected cluster1 (C), and cluster2 (D). E Top 10 Hub gene identified in PPI network of DEGs in PS-BCG using Cytohubba MCC based method.

Among 36 up-regulated critical network clusters in the case of the upregulated DEGs following *M. tuberculosis* infection, the highest MCODE score of 9.9. Among the 47 clusters identified in the case of the downregulated genes, the highest score was 81.3 **(Supplementary data file S5)**. **Fig 7A and B** depict the top two highly interconnected clusters. Functional analysis of these two clusters reveals extrinsic apoptotic signaling pathway, cellular response to IFN-, endothelial development, and antiviral response-related pathways in case of the cluster1 (**Fig 7C**), and nucleoside diphosphate metabolic process response to mitochondrial depolarisation and smooth muscle migration in case of cluster2 were major upregulated networks in case of *M. tuberculosis* infection (**Fig7D**). Additionally, by using Cytohubba-based analysis of the total upregulated genes following *M. tuberculosis* infection we identified *CXCL8, CXCL10, CCL5, TNF, CSF2, ICAM1, TLR3, IRF1, and NF-kB1A* as major upregulated hub genes amongst the complex protein interaction networks (**Fig7E**). Notably, hub genes identified for *M. tuberculosis* infections are distinct from the BCG group. Hub genes are central to biological mechanisms and represent potential signatures for an ongoing infection or disease state.

**Figure 7.**
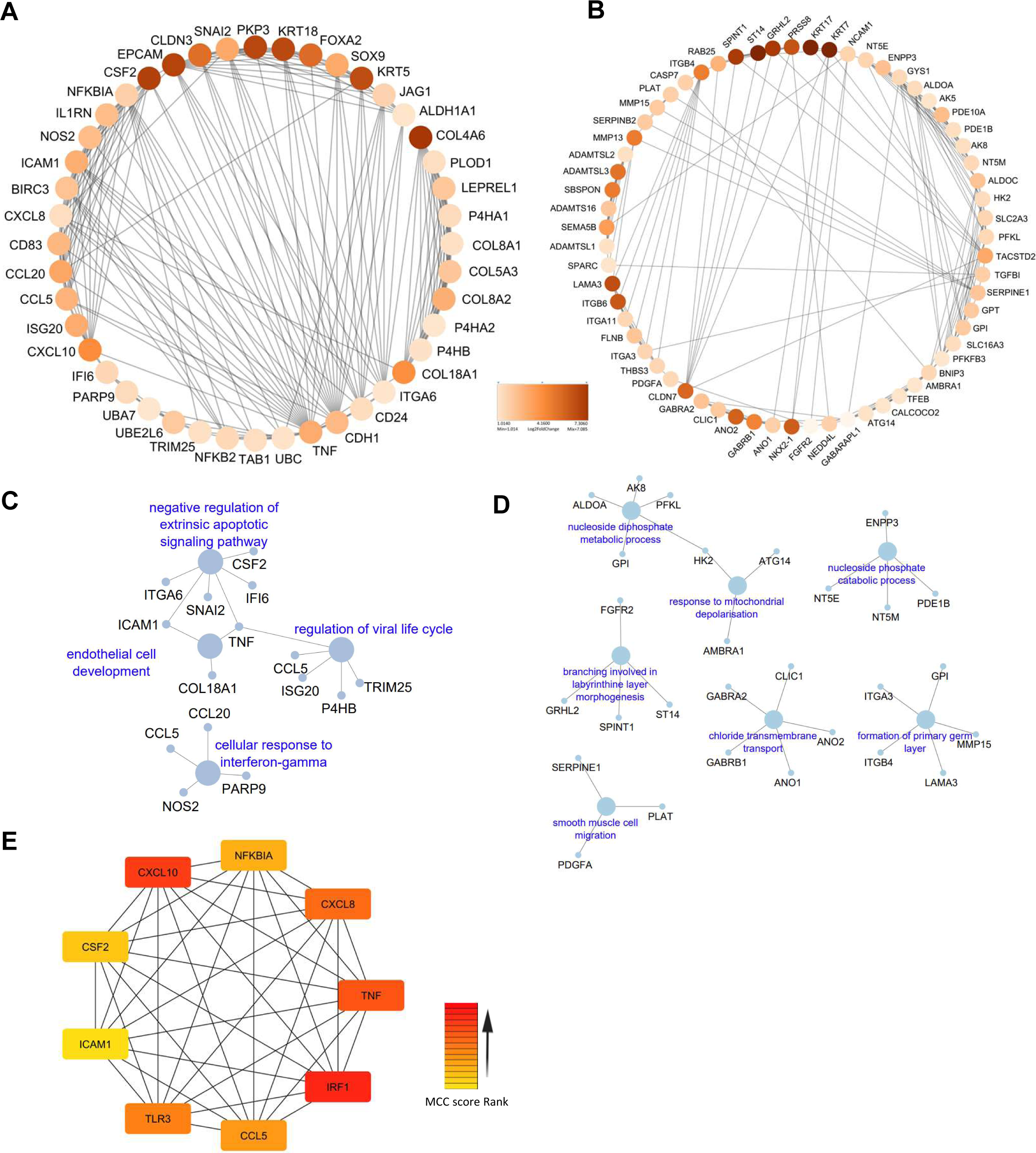
Network analysis of the up-regulated genes in *M. tuberculosis* infected pulmospheres. A, B Critical networks cluster1 (A) and cluster2 (B) were constructed using MCODE algorithm in Cytoscape. C, D ClueGO based GO of biological process, and immune related pathways in highly interconnected cluster1 (C) and cluster2 (D). E Top relevant Hub genes identified in PPI network of DEGs in PS-*M. tb* using Cytohubba MCC based method.

### Commonality and divergence of host responses to BCG and *M. tuberculosis* infection

Analysis of the DEGs identifies 388 common genes between BCG and *M. tuberculosis* groups, of which 307 showed a similar trend of either up or downregulation, and 81 genes exhibited inverse regulation (**Fig 8A**). Among the inversely regulated genes, 21 genes were upregulated and 59 were downregulated in *M. tuberculosis* compared to BCG infection (**Fig 8B**). GO analysis of the genes that are upregulated in case of BCG infection identified highly relevant immune response-related biological processes and pathways (leukocyte aggregation, type-2 immune responses, and other cytokine signaling pathways) in case of BCG infection (**Fig 8C**). Major genes involved in these immune response pathways were HCK, WNT5A, IL1R2, IL1RAP, IL33, TRIM62, EREG, MT2A, IL6, MMP1, MMP3, LRP8, and TNFRSF11B (**Fig 8B**). On the contrary, these genes were significantly downregulated in the case of *M. tuberculosis* infection indicating the possibility that *M. tuberculosis* infection subverts certain immune responses in its favor, and limits tissue repair and remodeling process. Notable genes that were significantly upregulated in the case of *M. tuberculosis* infection but downregulated in the case of BCG infection are RASSF5, CDH1, IGFBP2, TGFB3, and SPARC. Genes such as IGFBP2 and TGFB3 are involved in immune processes, while RASSF5 plays a role in apoptotic pathways, influencing host cell survival. Additionally, CDH1 and SPARC may modulate cell adhesion mechanisms and tissue remodeling processes, potentially impacting M. *tuberculosis* entry and dissemination within the host.

**Figure 8.**
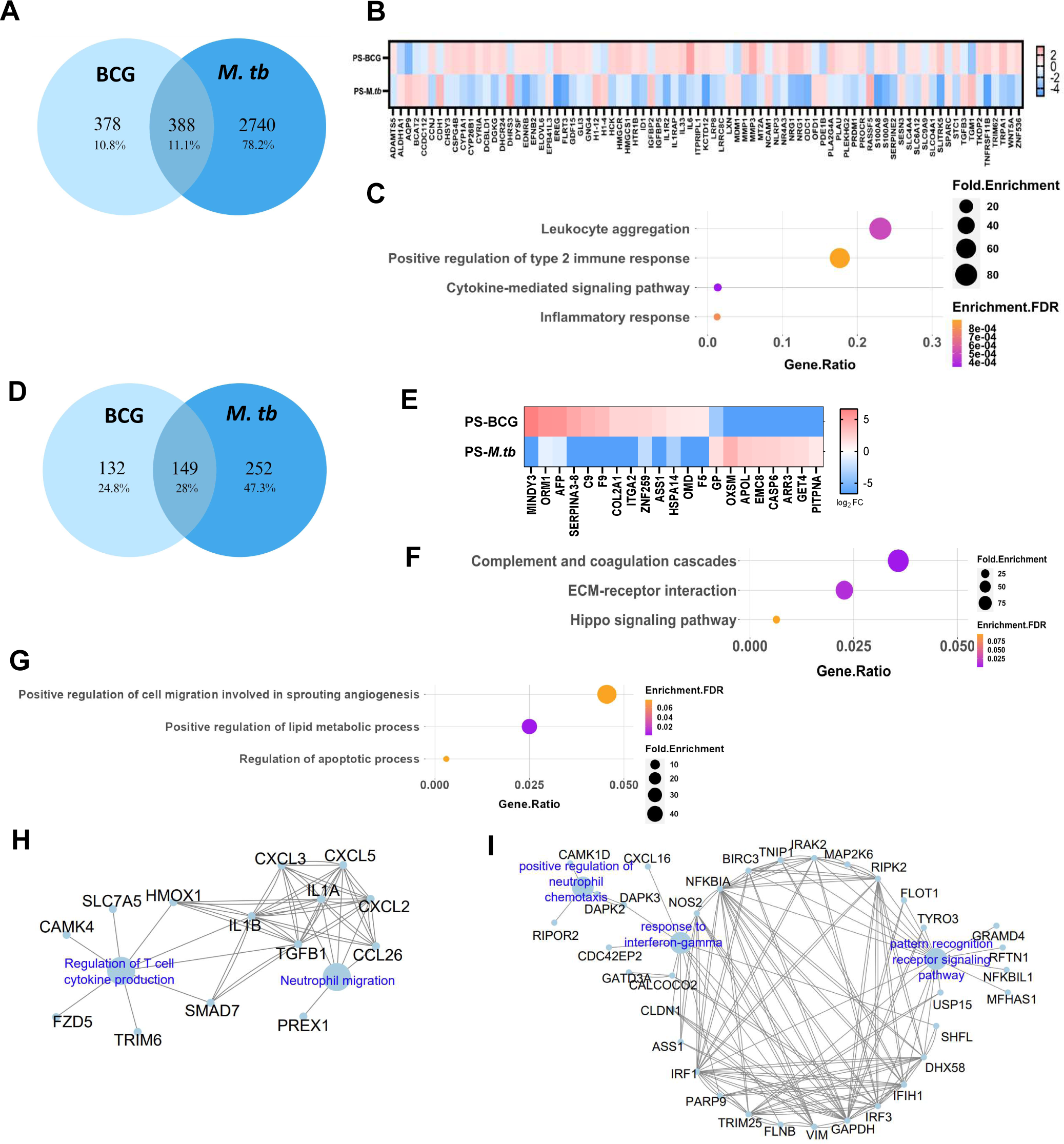
Divergence in the host responses to BCG and *M. tuberculosis* infection. A Venn diagram of DEGs shows common and unique genes upon BCG and *M.tb* infection of the pulmospheres. B Heat-map representing inversely regulated differentially expressed genes among common genes upon BCG and *M.tb* infection of the pulmospheres. C Top GO terms of common up-regulated genes in case of BCG infection and down regulated in *M.tb*. D Venn diagram of DEPs shows common and unique genes upon BCG and *M.tb* infection. E Heat-map representing the inversely regulated differentially expressed proteins upon BCG and *M.tb* infection. F Top GO terms of biological process for inversely regulated proteins that are upregulated in case if *M. tuberculosis* infection but down regulated in BCG infection. G Top GO terms of biological process up-regulated in case of BCG infection, but down regulated in case *M. tuberculosis* infection. H Immune network interaction that are up-regulated exclusively following BCG infection. I Immune network interaction upregulated exclusively upon *M. tuberculosis* infection.

Analysis of the DEPs identifies 149 common proteins between BCG and *M. tuberculosis* groups (**Fig 8D**), of which 128 showed a similar trend of either up or down-regulation, and 21 genes exhibited inverse regulation (**Fig 8E**). GO analysis on the 21 inversely regulated proteins highlights complement and coagulation cascade pathways, ECM-receptor interaction, and Hippo signalling pathway involving 13 genes as critical signalling pathways that are upregulated in case of *M. tuberculosis* infection but downregulated in case of BCG infection (**Fig 8F**). On the contrary, 8 genes belonging to cell migration and angiogenesis, lipid metabolism, and apoptosis were upregulated in the case of BCG infection, but downregulated in the case *M. tuberculosis* infection (**Fig 8G**).

Analysis of the unique DEGs in the case of BCG infection identifies 233 genes upregulated and 145 genes downregulated (**Fig 8A**). ClueGo-based immune responses related pathway analysis of the upregulated genes highlights neutrophil chemotaxis (CXCL3, CXCL2, CXCL5, CCl26, PREX1, IL1A, and IL1B) and regulation of T-cell cytokine production (TGFB, SMAD7, HMOX1, CAMK4, TRIM6, SLC7A5, and FZD5) as the significantly upregulated pathways (**Fig 8H**). Whereas, *M. tuberculosis* infection resulted in 1258 up and 1482 downregulated genes, and ClueGO analysis reveals enrichment of leukocyte activation, differentiation, migration, TLR-3 signaling, and response to IFN-γ pathways among the leading upregulated pathways (**Fig 8I**).

### Integrated analysis of transcriptome and proteome data identified potential biomarkers for early *M. tuberculosis* infection in the bovine lungs

Integration of transcriptomics and proteomics data highlighted key gene/protein expression signatures for *M. tuberculosis* and *M. bovis* BCG infection in the bovine pulmospheres. Sixteen genes/proteins in the case of BCG infection, 82 genes/proteins in the case of *M. tuberculosis* infection, and 59 proteins/genes uniquely expressed in the case of *M. tuberculosis* infection over BCG infection at both transcriptional and proteomic level (**Fig 9A**). Multi-gene correlation analysis reveals statistically significant positive correlation of 0.61 (*p=0.015*), 0.52 (*p=4.7e-07*), and 0.35 (*p=0.0062*), respectively for the genes commonly regulated at both transcriptional and proteomic level **(Fig 9B).** Next, by comparing the commonly upregulated genes/proteins from the three conditions discussed above (**Fig 9C, D, E**), we sort-listed 15 genes (COL17A1, CFB, APOA1, S100A2, SERPINE1, RBP4, TNFAIP8, ASS1, ITGA3, GYS1, ASL, MTHFD2, CLU, USP15, FDT1) as unique transcriptomic/proteomic marker of *M. tuberculosis* infection.

**Figure 9.**
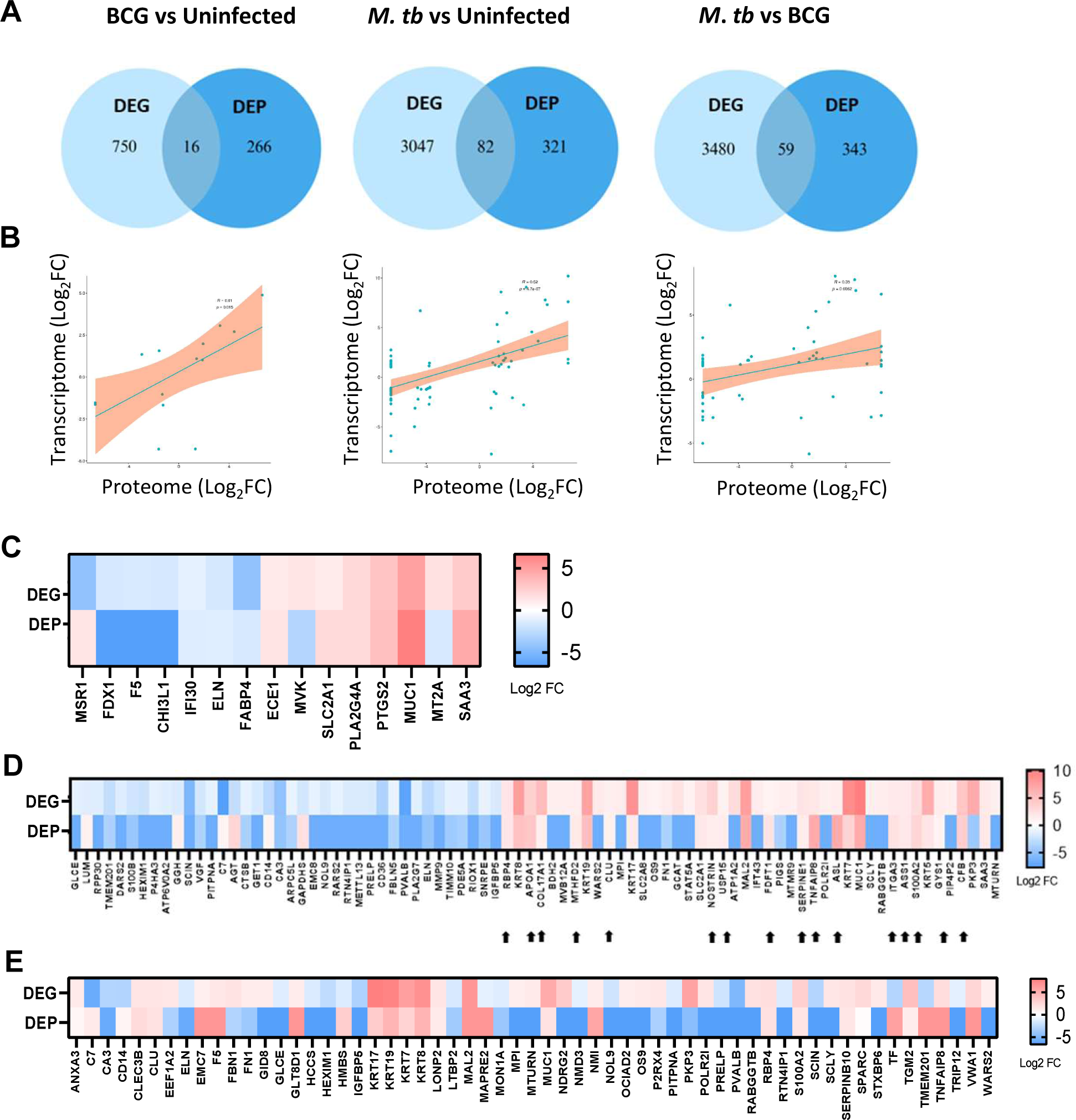
Integrated analyses of transcriptome and proteome data identify key markers of *M. tuberculosis* infection. A Venn diagram of DEGs and DEPs shows common and unique genes/proteins upon BCG and *M. tuberculosis* infection. B Correlation between Log_2_FC of transcriptome and proteome data for the common genes. P value < 0.05. C Heatmap representing the Log2 FC of DEGs and DEPs upon BCG vs uninfected. D Heatmap representing the Log2 FC of DEGs and DEPs upon *M. tuberculosis vs* uninfected. E Heatmap representing the Log2 FC of DEGs and DEPs upon *M. tuberculosis* infection vs BCG infection.

### *In silico* comparative analysis of potential *M. tuberculosis* infection biomarkers with published omics data

Nine hub genes identified from transcriptome data, and 15 upregulated genes, selected from the integrated analysis were chosen (24 genes) for further analysis for identifying potential markers of *M. tuberculosis* infection of the bovine lungs (**Fig 10A**). These genes/proteins could have significant potential as biomarkers for TB disease diagnosis, treatment monitoring, and the evaluation of vaccine-induced responses. We employed a robust in silico validation protocol to evaluate the cohort of identified genes. This involves a meticulous comparison with publicly available TB-transcriptome datasets (23 independent studies were included). The criteria for the selection of this dataset were described in detail in the materials and method section. **Fig 10B** illustrates the transcriptional status of selected 24 genes across all the data sets. Remarkably, 12 genes (IRF1, CXCL10, TNF, CXCL8, CCL5, ICAM1, COL17A1, CFB, SERPINE1, TNFAIP8, MTHFD2, and USP15) were found to be represented in more than 50% of the datasets. Finally, using a minimum cut-off Log2 fold induction of 0.585 (>1.5 fold in linear scale), and careful consideration of the nature of the bio-molecules, we shortlisted 7 genes/protein (IRF-1, CXCL10, CXCL8, CCL5, ICAM1, CFB, and SERPINE1) as potential *M. tuberculosis* early infection biomarker (**Fig 10C**).

**Fig. 10.**
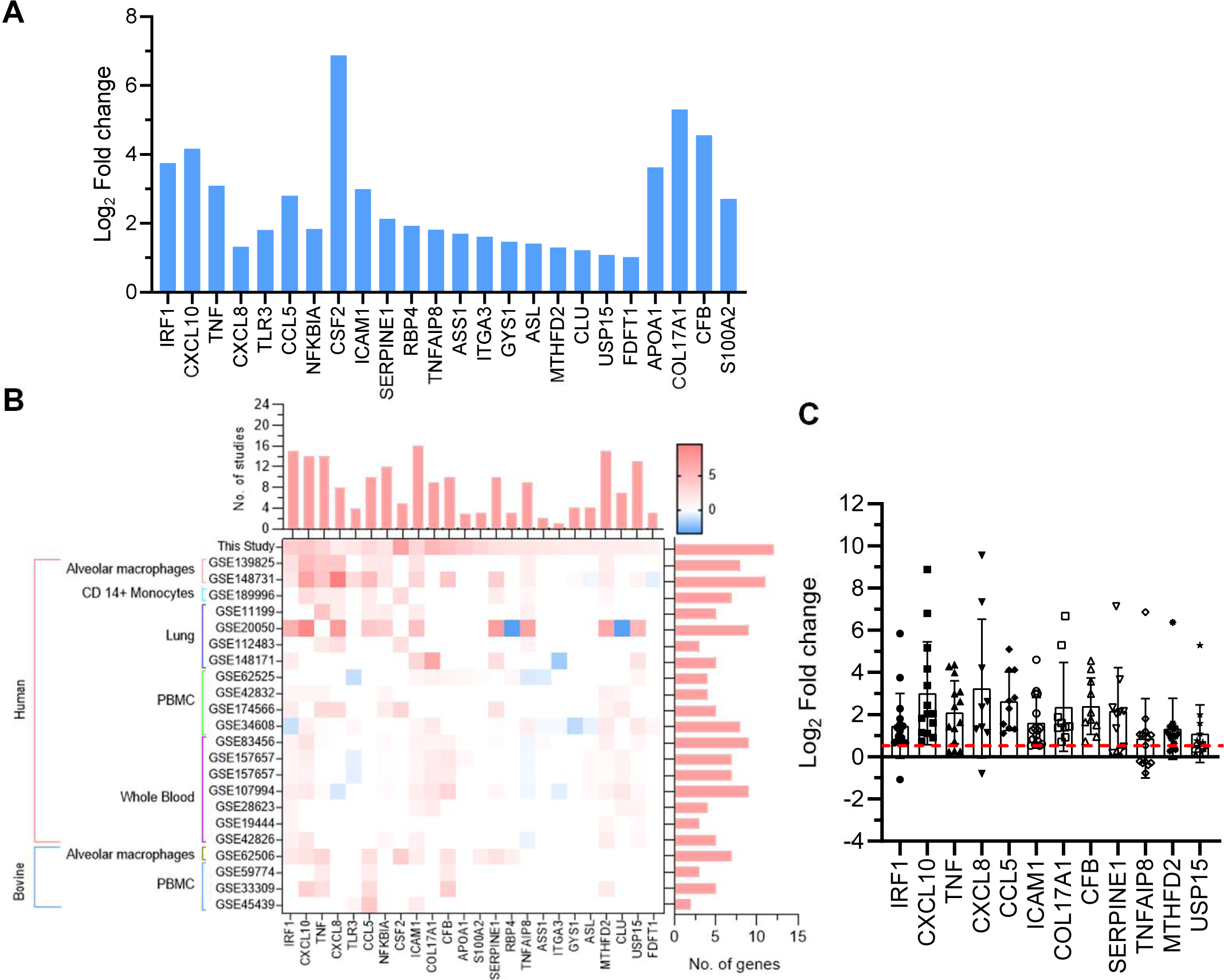
Potential biomarkers of early *M. tuberculosis* infection, and comparison with published omics data. A Bar diagram illustrates the Log2 fold change of 24 key genes selected in this study. B Heat-map represents the Log2 fold change of the key genes across selected similar omics studies, the bar diagram above the heat-map depicts the number of studies with up-regulation of the specific gene, and the bar diagram on the right side of the heatmap shows number of genes detected in a selected study. C The dot plot with mean and SD graph represents the Log2 fold change values of each gene across all the 23 selected studies. The red dashed line denotes the applied cut-off Log2 fold value of 0.585 (1.5-fold in linear scale).

## Discussion

In pursuit of a comprehensive understanding of bovine pulmonary TB, and the intricate host responses, this study undertakes the development of a robust and reproducible *ex vivo* disease model. The primary objective herein is to establish an innovative 3D pulmosphere model that faithfully recapitulates bovine TB and elucidates the fundamental signaling pathways and pivotal genes/proteins implicated in host-pathogen interaction during the early phase of TB infection within the bovine lungs. Notably, our 3D pulmosphere model fosters the co-culture of diverse cell types, encompassing macrophages, epithelial cells, fibroblasts, alveolar cells, and other immune cells intricately interconnected by the lung’s extracellular matrix components. This advancement stands in marked contrast to conventional 2D cell cultures, which frequently fail to replicate the intricate tissue architecture, cellular heterogeneity, and complexity of multicellular interactions. With its ability to model TB infection within the lungs with greater fidelity, the 3D pulmosphere model holds promise for deepening our insights into the bovine TB disease and facilitating the discovery of novel diagnostic biomarkers and therapeutic targets.

The ethical considerations surrounding research involving live animal models are noteworthy, and the *ex vivo* pulmosphere model offers a compelling solution. By utilizing bovine slaughterhouse-derived lung tissue, this model circumvents the ethical complexities associated with live animal experiments. Additionally, the bovine pulmosphere model possesses a scientific potential that extends to minimizing the need for live animals in research. In the context of bovine TB research, a notable challenge arises from the limited understanding of the initial stages of TB pathophysiology within the bovine lung due to the scarcity of studies, which is attributed in part to the intricate nature of conducting experimental infections in large animals within a bio-contained setting [14]. The pulmosphere model holds promise for reducing the number of animals required for basic research investigations on bovine TB and serves as an ethical and valuable substitute for large animal testing. An additional advantage of our pulmosphere model is its potential use as a cutting-edge in vitro model, offering significant promise for high-throughput screening of TB drugs. These three-dimensional miniaturized structures, mimicking the cellular organization and microenvironment of the lungs, provide a physiologically relevant platform for drug testing. High-throughput screening using bovine pulmospheres enables the simultaneous evaluation of multiple drug candidates, offering a cost-effective and time-efficient approach to identifying potential TB therapeutics. This can serve as an intermediate testing platform before laboratory animal-based drug screening, providing enhanced opportunities for the preclinical identification and evaluation of drug targets, as well as the identification of genes/proteins related to bacterial virulence. In summary, it represents an economical and ethically superior platform.

Our investigation employing the 3D pulmosphere uncovered distinctive immunological pathways, that distinguish between the responses induced by vaccine strain BCG and the virulent strain *M. tuberculosis*. Notably, *M. tuberculosis* infection precipitated a heightened divergence in gene expression and pathway regulation, accompanied by unique cellular signaling as compared to BCG infection. This divergence was evident in the heightened activity of the genes of complement and coagulation cascades and ECM receptor interactions within the 3D microenvironment of *M. tuberculosis*-infected pulmospheres, in contrast to BCG-infected pulmospheres. On the other hand, positive regulation of cell migration, lipid metabolism, and apoptotic process were the top up-regulated pathways following BCG infection, which corroborates previous findings [35, 36].

Further, a noteworthy finding: the heightened expression of interferon-stimulated genes (ISGs) and interferon regulatory factor (IRF) genes associated with both type-1 and type-2 interferon pathways during the initial phase of TB infection. While the important role of IFN-γ (type-2 interferon) in human TB is well known, the role of the Type-1 IFN pathway is conflicting, with some studies indicating its promotion of pathogen-favoring host responses and others suggesting the opposite [37–40]. However, its role in modulating TB disease in bovine hosts remains largely unexplored. In a precision-cut lung slice (PCLS) model, the Type-1 IFN pathway was found to be highly upregulated in the case of *M. bovis* infection compared to *M. tuberculosis* infection [9]. Our study leveraged the bovine 3D pulmosphere model to delve into the significance of this pivotal signaling pathway in the context of bovine TB. This approach holds promise for unearthing novel biomarkers for genomic selection initiatives, aiming to enhance cattle resistance to bovine TB. Furthermore, it offers a platform for developing innovative immuno-therapeutic strategies that can counteract the effects of the type-1 IFN pathway.

Intriguingly, our investigation highlights an up-regulation of the Th17 signaling pathway following exposure to BCG. Activation of the Rap1 pathway following BCG vaccination leads to heightened production of IL-17, a pivotal cytokine in combating mycobacterial infections [41]. Noteworthy evidence supports the induction of the IL-17 pathway following BCG vaccination, fostering protective immunity against *M. tuberculosis* in animal models [42]. Insights suggest BCG achieves this by stimulating IL-1β and IL-23 expression in those governing Th17 cell differentiation and expansion [43]. Additionally, the influence of BCG extends to dendritic cell activation, promoting Th17 differentiation and IL-17 production [44]. Notably, BCG induces a range of chemokines including CXCL1, CXCL2, CXCL5, and CCL20, recruiting Th17 cells to infection sites [45]. These Th17 cells in turn activate neutrophils and macrophages via IL-17, culminating in pathogen clearance [46]. Interestingly, *M. tuberculosis* infection also leads to IL-17 production, however, at a comparatively subdued level than BCG stimulation.

The interplay between BCG vaccination, *M. tuberculosis* infection, and tumor necrosis factor (TNF) signaling has been well documented in both animal models and humans [47]. Our innovative 3D bovine lung model, designed to mimic the lung environment, has revealed a robust TNF signaling pathway activation following infection with both *M. bovis* BCG and *M. tuberculosis*. Notably, the response was more pronounced in the case of the latter. TNF-α holds a pivotal role in the protective immune response against mycobacteria and is instrumental in the formation of granulomas, a hallmark of TB disease [47, 48]. BCG vaccination-induced TNF-α production proves indispensable for macrophage activation and mycobacterial eradication [49]. However, a delicate balance is necessary, as excessive TNF-α production can trigger tissue damage and inadvertently exacerbate disease progression [50]. In the context of *M. tuberculosis* infection, TNF-α assumes a critical role in orchestrating the recruitment and activation of immune cells, including macrophages and T cells, to the site of infection. *M. tuberculosis* capitalizes on the TNF pathway to its benefit, inducing the expression of TNF receptor 2 (TNFR2) on macrophages, thereby promoting bacterial survival and growth [51]. This multifaceted involvement of TNF signaling underscores its intricate role in both defense and subversion strategies during mycobacterial infections, and the 3D-pulmosphere model highlights its potential as a useful model to study TNF signalling in TB and therapeutic interventions targeting this critical pathway.

The heightened expression of the Advanced Glycation End-products (AGE) and Receptor for AGE (RAGE) pathway (AGE-RAGE) has been observed in *M. tuberculosis*-infected pulmospheres, highlighting its relevance in TB pathogenesis [52, 53]. This pathway known for its significance in various disease contexts, plays a crucial role in TB as well. The formation of AGEs through non-enzymatic glycation of proteins and lipids is linked to inflammation and oxidative stress in diverse tissues. The Receptor for AGEs (RAGE), present on immune cells, becomes activated upon interaction with AGEs, initiating signaling cascades including NF-kB and MAPKs, and subsequently leading to the production of pro-inflammatory cytokines and chemokines [54]. In the TB context, the AGE-RAGE pathway emerges as a key contributor to lung damage, fibrosis, and granuloma formation [55]. Intriguingly, BCG vaccination is implicated in the modulation of this pathway. Studies reveal a reduction in AGE accumulation and RAGE expression in the lungs of BCG-vaccinated mice subsequently infected with *M. tuberculosis*, offering insights into the vaccine’s potential impact on this detrimental pathway [56]. Our finding underscores the AGE-RAGE pathway’s role in bovine TB pathogenesis.

Significant perturbations in signaling pathways also encompass the Peroxisome proliferator-activated receptor (PPAR) pathway, notably upregulated in both BCG and *M. tuberculosis*-infected pulmospheres in this study. This pathway assumes a pivotal role in regulating lipid metabolism and immune responses [57]. In the context of TB, PPAR signaling emerges as a critical modulator of host immune responses to *M. tuberculosis* infection [58]. The research underscores the capability of PPAR-gamma agonists to bolster host defense against *M. tuberculosis* by fostering antimicrobial peptide production and augmenting macrophage phagocytosis [59]. Conversely, PPAR-delta exhibits an inhibitory impact on host defense by tempering pro-inflammatory cytokine release and bolstering bacterial survival within macrophages. Of note, PPAR-gamma agonists can heighten BCG vaccine immunogenicity by promoting dendritic cell activation and Th1 cytokine production [60]. Furthermore, insights suggest BCG vaccination can activate the PPAR signaling pathway, subsequently influencing the expression of genes pivotal to lipid metabolism and immune responses [35].

Our *in silico* comparative analysis of published OMICS data with our experimental findings identifies 7 genes/proteins (IRF-1, CXCL10, CXCL8, CCL5, ICAM1, CFB, and SERPINE1) as potential early infection biomarkers for *M. tuberculosis* infection in the bovine pulmospheres. IRF-1 is a transcription factor that contributes to the activation of macrophages, and promotes the expression of several pro-inflammatory cytokines including IL-6, TNF-α, and IL-8 [61–63]. IRF1 is known to exert its anti-*M. tuberculosis* effect via suppressing the (mTOR)/p70 S6 kinase (p70 S6K) cascade [61], and IRF-1 has also been proposed as biomarker for active TB in humans [64]. In the case of TB, CXCL10 is induced by IFN-γ and has been associated with the recruitment of T cells to the lungs [65]. A plethora of studies reported elevated levels of CXCL10 in TB patients, indicating its potential role in the immune response to *M. tuberculosis* infection [66]. Our bovine pulmosphere model-based findings of CXCL10 as an early TB biomarker resonate with a prior investigation that identified CXCL10 as a potential cell-mediated immuno-biomarker for bovine TB diagnosis [21]. Shortlisting of CCL5 (RANTES, regulated upon activation, normal T cell expressed and secreted) and CXCL8 (IL-8) as potential TB infection markers augur well with the two previous markers. In TB, both CCL5 and CXCL8 is produced by various cell types, including macrophages and epithelial cells, however, CXCL8 is primarily involved in the recruitment and activation of neutrophils at the site of infection whereas CCL5 is involved in the recruitment of T cells, monocytes, and eosinophils [67, 68]. Notably, CCL5 and CXCL8 chemokines have been proposed as potential biomarkers of active TB and latent TB, respectively [69, 70]. ICAM1 (Intercellular Adhesion Molecule 1) is a cell adhesion molecule that facilitates the interaction between immune cells, such as T-cells and endothelial cells, and contributes to the recruitment of immune cells to the site of infection during TB [71]. Several studies proposed ICAM1 as a potential serum protein biomarker associated with TB [72, 73]. CFB (Complement Factor B) is part of the alternative pathway of the complement system and contributes to the formation of the C3 convertase complex and subsequent opsonization [74]. Evidence suggest that as a part of the acute phase response, the complement system may influence the inflammatory response during TB via activation of complement components, including CFB, leading to the generation of anaphylatoxins and the recruitment of immune cells to the site of infection thereby contributing to bacterial opsonization and phagocytosis of *M. tuberculosis* by immune cells highlighting the role of CFB in the innate immune response to tuberculosis [75]. Previously, elevated C1q level was considered a potential biomarker for active TB compared to LTBI, and CFB was proposed as a novel biomarker candidate for pancreatic ductal adenocarcinoma, this is the first instance, we found CFB could be a potential marker for early *M. tuberculosis* infection [76, 77]. Finally, SERPINE1 (Serpin Family E Member 1), also known as plasminogen activator inhibitor-1 (PAI-1), is involved in the regulation of the fibrinolysis pathway [78]. In TB, PAI-1 has been associated with the modulation of the inflammatory process to balance between inflammation and tissue repair [79], and its levels were found to be elevated in TB patients [80, 81]. Due to the similarity of high PAI-1 levels in the case of other pathological conditions and lung infections, it has not been exclusively proposed as an active TB biomarker in humans [79, 82–84]. While, PAI-1 has never been proposed as a Bovine TB biomarker, bovine alveolar macrophage showed transcriptional activation of PAI-1 upon *M. bovis* infection [85]. The robustness of the in-silico analysis and the consistent representation of these biomarkers across diverse datasets enhance the credibility and potential clinical utility of the identified early infection biomarkers. However, further experimental validation is warranted to confirm their diagnostic efficacy and clinical applicability.

While our 3D pulmosphere model offers versatile utility and suitability, it is important to acknowledge certain limitations that warrant further improvement and refinement. Creating and maintaining the 3D pulmosphere model presents greater challenges compared to traditional 2D cell culture, potentially constraining its adoption in certain laboratories [24]. Furthermore, the model’s ability to replicate the intricate *in vivo* environment is not exhaustive, as it lacks the natural recruitment of immune cells from peripheral sites to the infection foci post-infection. Addressing this limitation could involve integrating relevant peripheral blood mononuclear cell (PBMC)-derived immune cells with lung-derived cells for pulmosphere generation. Additionally, it’s noteworthy that this study focused solely on host responses at the 24-hour post-infection mark; investigating responses during different phases of TB infection and granuloma formation using this model could enhance our comprehension of bovine TB pathogenesis and consequently refine intervention strategies. Further investigations must be conducted to confirm the functional significance of the identified genes and pathways in the pathogenesis of bovine TB. Additionally, as it is well-known that the virulence of *M. tuberculosis* in bovine is relatively lower compared to *M. bovis* [86], the use of *M. tuberculosis* as the virulent tubercle bacilli in this study may not represent the true bovine pulmonary response to *M. bovis* infection. Our future studies using the 3D-pulmosphere model to include *M. tuberculosis, M. bovis* as well as the rapidly emerging leading MTBC pathogen *M. orygis* for a comparative investigation of the host response to bovine TB would address this limitation [87]. Notwithstanding these considerations, our findings offer valuable insights into the biological pathways and molecular mechanisms governing the bovine host’s response to BCG and *M. tuberculosis* infections at the primary site of infection.

In summary, this is the first report demonstrating the development of a superior *ex vivo* 3D pulmosphere bovine TB model. The combination of diverse cell types and increased ECM-related protein expression in 3D pulmospheres emphasizes their physiological relevance confirming the value of 3D pulmospheres for studying host-pathogen interactions beyond 2D cultures. Transcriptomic and proteomic analyses provide a comprehensive view of the early host response to *M. tuberculosis* in bovine 3D pulmospheres, revealing species-specific responses, hub genes, and functional clusters. These findings offer new insights into the molecular mechanisms governing host-pathogen interactions during TB in bovines, with potential implications for the discovery of disease biomarkers, host-directed therapies, and enhancing our understanding of TB pathogenesis, and ultimately aiding in the discovery of improved prevention and management strategies.

## Materials and methods

### Lung tissue and single-cell suspension preparation

Bovine lung tissue samples were collected from the approved abattoir and transported in ice-cold phosphate-buffered saline (PBS) containing antibiotic-antimycotic to the laboratory. After thorough washing in PBS, the tissue was chopped into small pieces of approximately 2mm X 2mm size and kept in 5 ml dPBS containing liberase enzyme (5mg in 1ml/1 gm of tissue) and was incubated at 37 °C in a 50 ml falcon tube for 40 minutes with intermittent agitation (every 10 mins). Following liberase treatment, 5ml of DMEM complete media containing 1x HPES, 10% FBS was added and shaken vigorously to disassociate the tissue. The dissociated cells were filtered through a 100-μm filter and pelleted at 1500 rpm for 5 mins at room temperature (RT). Subsequently, cells were washed with dPBS, and red blood cell (RBC) lysis was performed using RBC Lysis Buffer^TM^. Cells were then washed twice with dPBS, followed by DMEM complete media, and finally resuspended in DMEM-F12 complete media (consisting of 10% FBS, 1X Antibiotic-Antimycotic solution). Cells were seeded into a T-25 flask grown for 7 days, and subsequently used for pulmosphere preparation.

### Preparation of poly-HEMA–coated plates

For a generation of 3D pulmospheres, we followed the protocol described previously [88]. Briefly, Poly 2-hydroxyethyl methacrylate (poly-HEMA) stock solution was prepared (120 mg/ml) in 95% ethanol by dissolving the crystals at 65°C on a heat block for 10-12 hours. The working solution of poly-HEMA (5 mg/ml) was prepared vortexed briefly for 30 seconds and added into 96-well U-bottom plates (160-μl/well). Plates were left at RT in a sterile laminar hood for 24 hours until the plates were fully dry. These coated plates were used for the preparation of the pulmospheres.

### Preparation of the pulmosphere

Seven days cultured primary lung cells were detached from the T-25 flask using trypsin (0.5%) and neutralized with 5 ml DMEM complete media. Cells were collected by centrifugation at 300g for 5 mins at RT. The cell pellet was resuspended in a 2 ml complete DMEM medium, and cell viability and live cell density were measured via the trypan blue exclusion method. Subsequently, 8,000–10,000 cells were added per well in the poly-HEMA–coated 96-well U-bottom plates. The plate was centrifuged at 200g at RT for 1 min and was incubated for 24 hours at 37°C and 5% CO_2_ in a TC incubator resulting in the formation of spheroids. These multicellular 3D-spheroids derived from the primary lung cells were termed “pulmospheres”. The day wise growth of pulmosphere was measured using image J and motic image plus 3.0 software.

### Mycobacterial cultures

The *Mycobacterium bovis* BCG (Danish 1331 strain) and *M. tuberculosis* (H37Rv strain) were cultured in Middle Brook (MB) 7H9 broth containing 1x Oleic Acid Dextrose Catalase (OADC) supplement and .02% Tween 80 following standard method [89]. Bacteria from the mid-log phase broth culture were used for the preparation of glycerol stocks and stored at -80°C for subsequent use. For the *in vitro* infection, fresh bacterial cultures were grown from the glycerol stock to the mid-log phase, washed thoroughly with 1X PBS, and resuspended in DMEM incomplete media (without FBS and antibiotic-antimycotic mix) for use in the infection experiment.

### Bovine Pulmosphere TB Infection Model

To establish the 3D-pulmosphere *M. tuberculosis* infection model, cultured primary lung cells from passage number 0 or 1 were infected with the tubercle bacilli from logarithmic phase bacterial culture at a pre-calibrated multiplicity of infection (MOI) of 1:10 (cell: bacteria). Following three hours of infection cells were thoroughly washed to remove the extracellular bacteria. Cells were then detached from the 2D culture using trypsin (0.5%) and single cell suspension of infected mixed cells was used for the formation of *M. tuberculosis*-infected 3D pulmosphere using the same method described in the above sub-section ’Preparation of the pulmosphere’. Additionally, *M. bovis* BCG was also used to generate BCG-infected 3D-pulmopheres following the same protocol.

### Sample preparation for mass spectrometry

For the preparation of each sample for mass spectrometry (MS) analysis, ten pulmospheres from each treatment group were pooled containing approximately 1 × 10^5^ cells, and lysis was performed using lysis buffer containing 2% Sodium deoxycholate (SDC) in 50 mM Tris HCL (PH 7.5). Protein concentrations were measured by Bradford assay. The protein samples were treated with dithiothreitol at 57°C for 1 hr to reduce disulfide bonds and subsequently alkylated by iodoacetamide treatment for 1hr in the dark at RT. Then samples were subjected to trypsin digestion overnight at 37°C at a ratio of 1:20 (trypsin: protein) followed by SDC separation via formic acid (25%) treatment. Digested protein samples were then filtered through a 10KDa membrane filter to remove undigested proteins followed by salt removal using the C18 column and samples were vacuum dried. Finally, the dried samples were reconstituted in 0.3% formic acid, quantified using a spectrophotometer, and 1 µg of peptide samples were subjected to tandem MS-MS analysis.

### Mass spectrometry data acquisition and data analysis

The Ultimate 3000 RSLCnano system coupled with the high-resolution Q Exactive HF mass spectrometer (Thermo Fisher) was used. The full MS1 scans were set with a resolution of 60,000 ions, an automatic gaining control (AGC) target value of 1 × 10^6^, and an acquisition range of 375– 1600 m/z with a 60 ms maximum injection time. For the fragmentation, the top 25 precursors were selected. For the MS2 acquisition, a resolution of 15,000 ions, a target AGC value of 1 × 10^5^, with a maximum injection time of 100 ms, an isolation window of 1.3 m/z, and a fixed first mass at 100 m/z were specified. Peptide elution was performed with a nonlinear gradient flow of 0.300 µL/min during 180 min using 5% of solvent B (80% acetonitrile + 0.1% formic acid) and 95% of solvent A (0.1% formic acid). The data was analyzed using proteome discoverer v2.5 (Thermo Fisher).

### RNA extraction from the pulmosphere

RNA isolation was performed using a combination of trizol (sigma) and RNeasy Mini plus Kit (Qiagen). Briefly, 1ml trizol was added to the pooled pulmospheres samples, vortex vigorously and 200 μL chloroform was added. Samples were then shaken for 15 seconds, incubated for 3 minutes at RT, and then centrifuged at 12,000g for 15 minutes at 4°C. The upper aqueous layer was separated and proceeded further for RNA extraction using RNeasy Mini kit following manufacturer’s instructions. Total RNA was eluted with 30 μL RNase-free H_2_O and stored at – 80°C. the RNA concentration and purity was checked using Nanodrop 1000 (Thermo Fisher).

### Whole transcriptome sequencing and analysis

RNA quantity is checked with Qubit fluorometer (Thermofisher #Q33238) using RNA HS assay kit (Thermofisher #Q32851) following manufacturer’s protocol, and subsequently, RIN values were estimated in a tapestation 4150 using HS RNA screen tape (Thermo Fisher). The library preparation was carried out using TruSeq® Stranded Total RNA kit (Illumina #15032618, Illumina #20020596). Final libraries were quantified using Qubit 4.0 fluorometer (Thermofisher #Q33238) using a DNA HS assay kit (Thermofisher #Q32851) following the manufacturer’s protocol. To identify the insert size of the library, we queried it on Tapestation 4150 (Agilent) utilizing highly sensitive D1000 screentape (Agilent # 5067-5582) following manufacturers’ protocol. Quality assessment of the raw fastq reads of the sample was performed using FastQC v.0.11.9 (default parameters) [90]. The raw fastq reads were preprocessed using Fastp v.0.20.1 (parameters: -- trim_front1 9 --trim_front2 9 --length_required 50 –correction --trim_poly_g -- qualified_quality_phred 30) [91], followed by quality re-assessment using FastQC and summarization using MultiQC [92]. The processed reads were aligned to the STAR indexed Bos taurus ARS-UCD1.2 genome using STAR aligner v 2.7.9a (parameters:’--outSAMtype’ BAM SortedByCoordinate, ‘--outSAMunmapped’ Within, ‘--quantMode’, TranscriptomeSAM, ’--outSAMattributes’ Standard) *Bos Taurus* (ARS-UCD1.2) [93]. The rRNA features were removed from the GTF file of Bos taurus genome ARS-UCD1.2. The alignment file (sorted BAM) from individual samples was quantified using featureCounts v. 0.46.1 based on the rRNA-filtered GTF file to obtain gene counts [94]. These gene counts were used as inputs to DESeq2 for differential expression estimation (parameters: threshold of statistical significance --alpha 0.05; p-value adjustment method: BH) [95]. For Gene Ontology (GO) and Kyoto Encyclopedia of Genes and Genomes (KEGG) Pathway annotation, the up and down-regulated gene IDs were extracted from the DESeq2 result files and subjected to bioDBnet [96]. ShinyGO 0.77 was used for functional enrichment analysis, and cross-verified with g:Profiler. Venn diagrams were produced using Venny 2.1. Heatmaps were generated following hierarchical clustering, and discriminating variables between comparison groups were identified using a false discovery rate of p < 0.005 or q < 0.2.

### Network analysis

Up-regulated and down-regulated genes from proteome and transcriptome were used to construct protein-protein interaction network (PPI) using Online Database STRING. Full string network types having both physical and functional association with medium confidence 0.4 were kept as default settings. The resultant network was imported to Cytoscape software (3.9.1) to analyze the PPI network [97]. Highly connected clusters were identified using MCODE clustering method. The highest MCODE score in the network was subjected to GO enrichment analysis using ClueGO setting a two-sided hypergeometric test with Bonferroni step-down corrected p-value ≤ 0.05, and kappa score ≥ 0.4 as statistical parameters. The maximal clique centrality (MCC) method was applied to identify the important hub genes based on the MCC Score [98].

### *In silico* comparative analysis for validation of *M. tuberculosis* infection signature

We implemented a rigorous validation protocol to assess a set of identified genes that potentially serve as key regulators in bovine tuberculosis (TB) conditions. To validate these key genes, we compared them against publicly available TB infection transcriptome datasets. We specifically focused on multiple TB disease cohort studies accessible through the NCBI Gene Expression Omnibus (GEO) database (https://www.ncbi.nlm.nih.gov/geo/). The list of selected studies, along with their GEO accession numbers, is provided in **Supplementary data file S6**. Our selection criteria were confined to two species: Bovine and Human. We used search terms such as "Bovine tuberculosis host transcriptomic", "Bovine tuberculosis RNA seq", "Tuberculosis lung transcriptome", "Tuberculosis blood signature", "Tuberculosis lung signature", "Tuberculosis host RNA seq", "Tuberculosis PBMC gene expression", "Tuberculosis whole blood gene expression", "Tuberculosis lung gene expression", and "Tuberculosis sputum". These terms encompassed both microarray and high-throughput sequencing data types. The dataset inclusion criteria encompassed studies involving *M. tuberculosis* and *M. bovis* infections specifically in peripheral blood mononuclear cells (PBMC), whole blood, lung, and alveolar macrophages (AM). In broad terms, the analyses centered around comparing groups infected with *M. tuberculosis* and *M. bovis* with their respective healthy control groups. Based on the above inclusion factors 22 studies were considered for further comparison. The identification of DEGs was performed using GEO2R, integrated with the limma package, considering genes with statistically significant differences between pairwise groups (adjusted P value < 0.05, FDR < 0.05).

### Statistical Analysis

The statistical analysis and preparation of graphs were performed using GraphPad Prism 9 software (GraphPad, CA, USA). Data are presented as mean ± sd. Statistical analysis included an unpaired two-tailed t-test for comparison between two groups unless otherwise specified in the figure captions. A 95% confidence interval or 0.05 threshold for significance (p < 0.05) was used in all statistical tests. For correlation analysis, GraphBio was used [99].

### Databases used for analysis

Several publicly available databases were used to aid in the in-depth analysis of cell types, networks, and pathways based on the transcriptome and proteome data. Some of the major databases include Matrisome DB for ECM atlas [100]. All the databases and software information are provided in the **Supplementary data file S7.**

## Ethics statement

All experiments were reviewed and approved by the Institutional Biological Safety Committee (IBSC, approval no. IBSC/2018/NIAB/BD/001) of the National Institute of Animal Biotechnology, Hyderabad, and animal experiments were approved by the Animal Ethics Committee of the West Bengal University of Animal and Fishery Sciences, Kolkata, India (Approval No. IAEC/22-B, CPCSEA Reg. No.763/GO/Re/SL/03/CPCSEA), and the Committee for the Purpose of Control and Supervision on Experiments on Animals, India. All procedures were performed in accordance with the relevant guidelines and regulations laid down by Govt. of India.

## Data availability

All transcriptome and proteome data are available in the NCBI GEO accession number GSE246765, and in ProteomeXchange consortium via PRIDE [96] partner repository with the database identifier PXD046641, respectively. All other data supporting the findings of this study are available from the corresponding author upon reasonable request.

## Author Contributions

Conceived the project: BD. Conceived and designed the experiments: VB, BD. Performed the experiments: VB, RK, SG, HKM, US, BD. Analyzed the data: VB, RK, MRP, US, BD. Contributed reagents/ materials/analysis tools/ facility: BD, US. Wrote the paper: VB, RK, BD. Provided overall supervision throughout the study: BD.

## Acknowledgment

We are thankful to Dilna S V, and Nilanjana Ganguli for technical help with the LC-MS analysis. We acknowledge Prof. Sharmistha Banerjee, Co-ordinator of the UoH-NIAB BSL3 facility, University of Hyderabad, India, and other technical staff for supporting the BSL3-based experiments.

## Statements and Declarations

### Supplementary information

Please see the supplementary information file.

