## Supplementary figures for "A bovine pulmosphere model and multiomics analyses identify a signature of early host response to *Mycobacterium tuberculosis* infection"

**Title:**

### Supplementary Figure S1

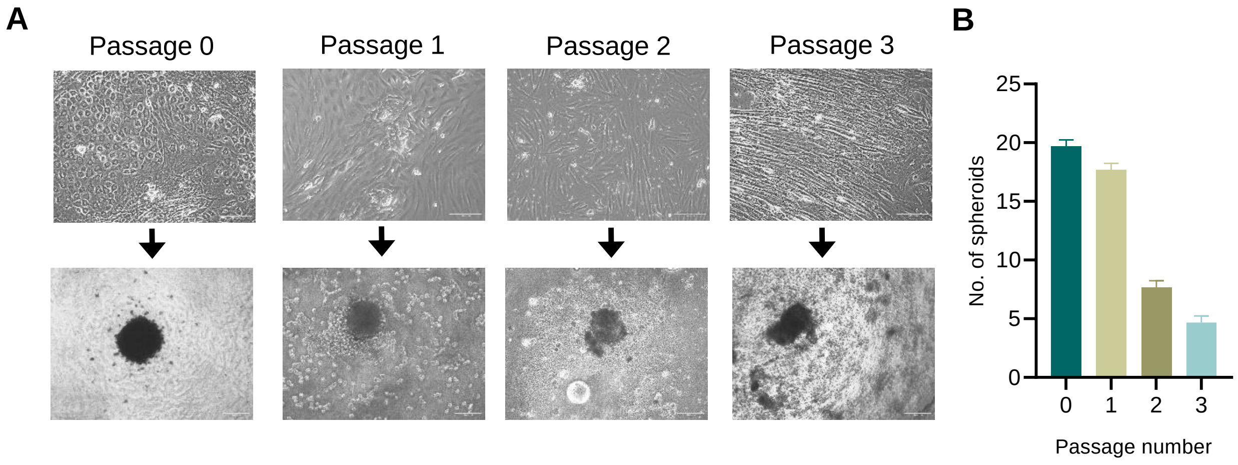


**Supplementary Figure S1. Influence of passage number on 3D pulmosphere formation.**

A Representative images of bovine primary lung cells monolayer culture at successive passages, and corresponding pulmospheres derived from them. Scale bar 200 µm for 10X images and 400 µm for 4X images.

B Bar graph depicts the average number of well-formed pulmospheres derived from different passages of primary lung cells. Data represented in mean ± sd.

### Supplementary Figure S2

**A**

#### Proteomics sample preparation and LC-MS workflow

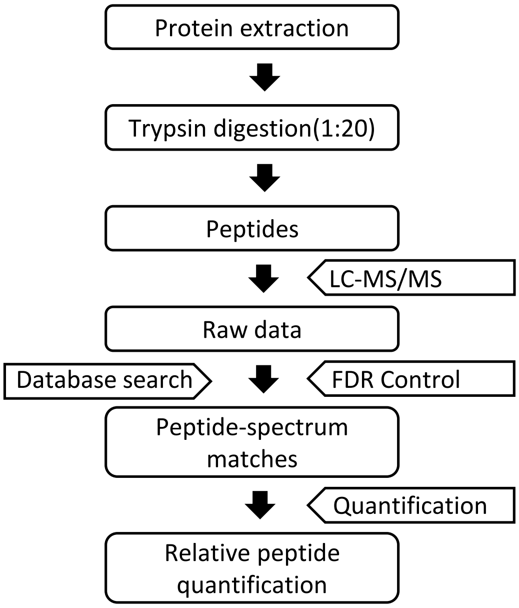


**B**

#### Proteomics data analysis workflow

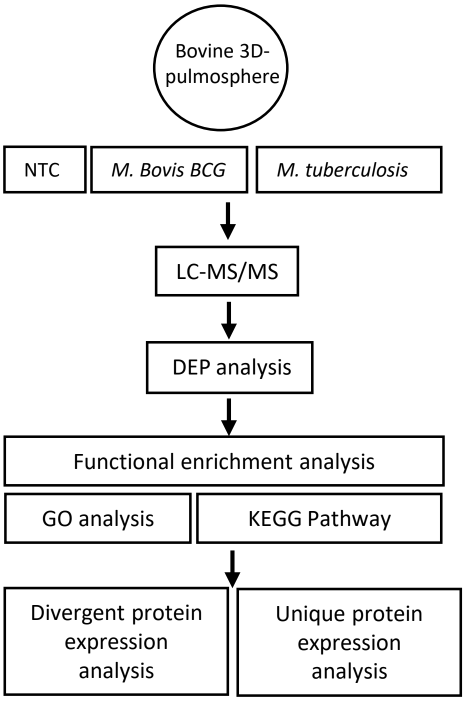


**Supplementary Figure S2. Workflow for the LC-MS sample preparation and data analysis**.

A Schematic representation of the protocol for the sample preparation of bovine 3D pulmosphere for LC-MS/MS analysis.

B Analysis workflow for LC-MS/MS data of 3D pulmosphere with or without infection.

### Supplementary Figure S3

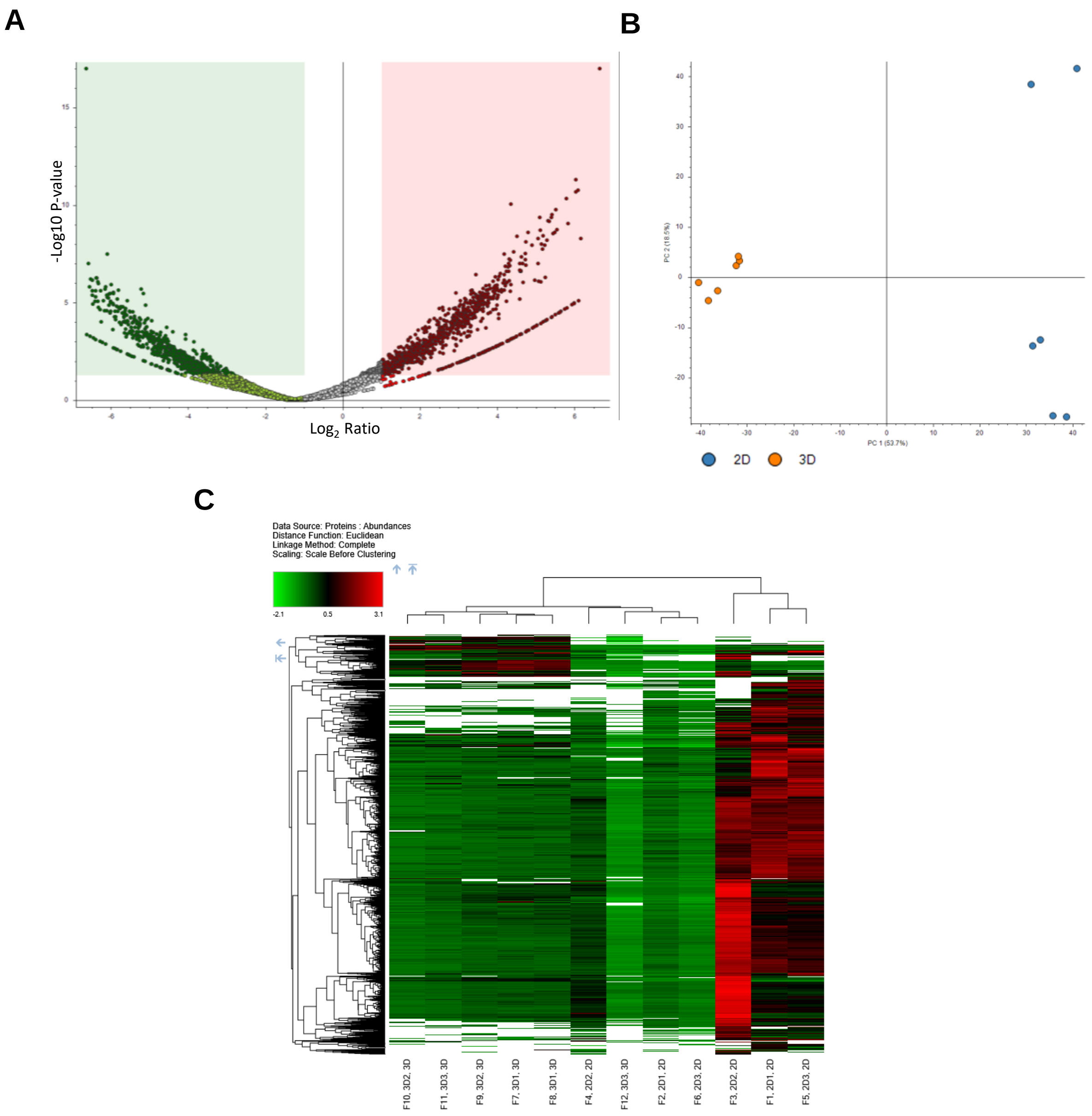


**Supplementary Figure S3. Comparative proteomic analysis of 3D pulmosphere vs 2D lung monolayer cell culture**.

A Volcano plot depicts up and down-regulated proteins in 3D pulmosphere based on the $Log_2$fold change in comparison with 2D monolayer culture. Red and Green dots indicate up and down-regulated proteins, respectively; FDR cutoff is <0.05 and $Log_2$1.

B Principle component analysis (PCA) of 3D pulmosphere (n=3) and 2D lung cell monolayer culture (n=3). PC1 and PC2 represent the % of variance between and among the sample groups.

C Hierarchical clustering analysis of peptide abundances in 3D vs 2D samples.

### Supplementary Figure S4

#### A Transcriptome sequencing workflow

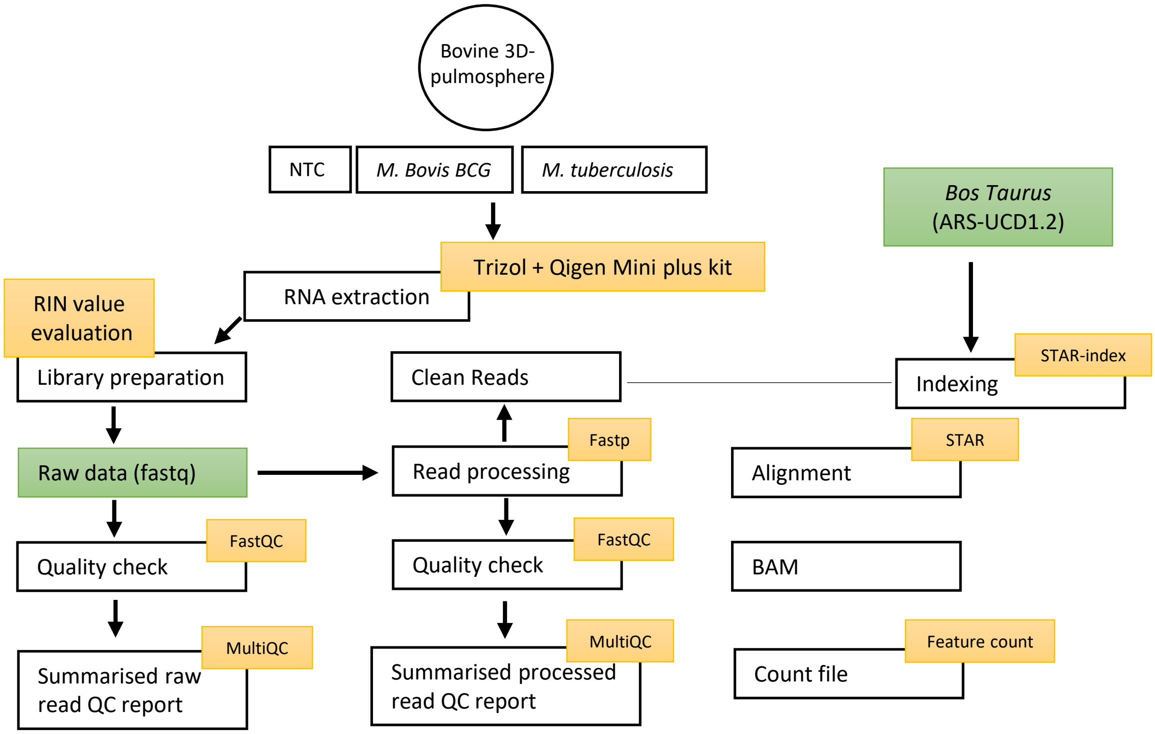


#### B Transcriptome data analysis workflow

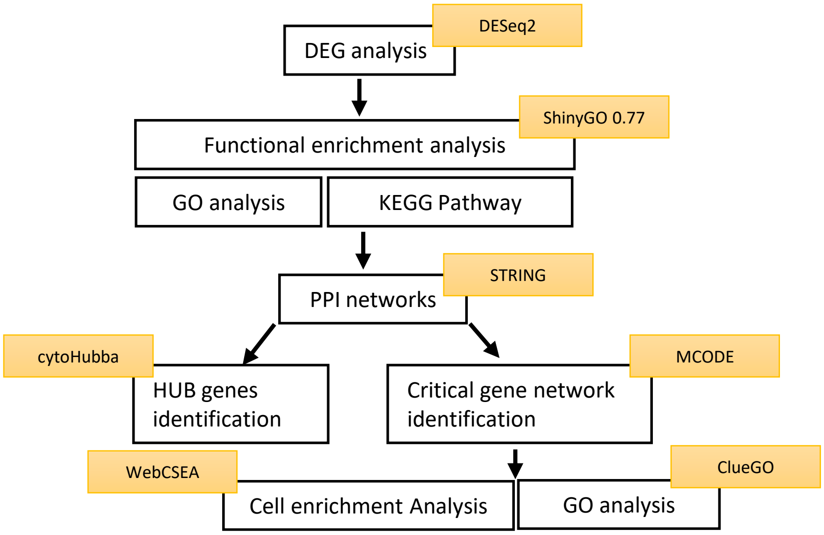


**Supplementary Figure S4. RNA seq workflow and downstream analysis.**
A Flowchart of the protocol for the RNA sample preparation, library construction and data processing of bovine 3D pulmosphere based RNA seq experiment.
B Analysis work flow of RNA Seq data of 3D pulmosphere with or without infection.

### Supplementary Figure S5

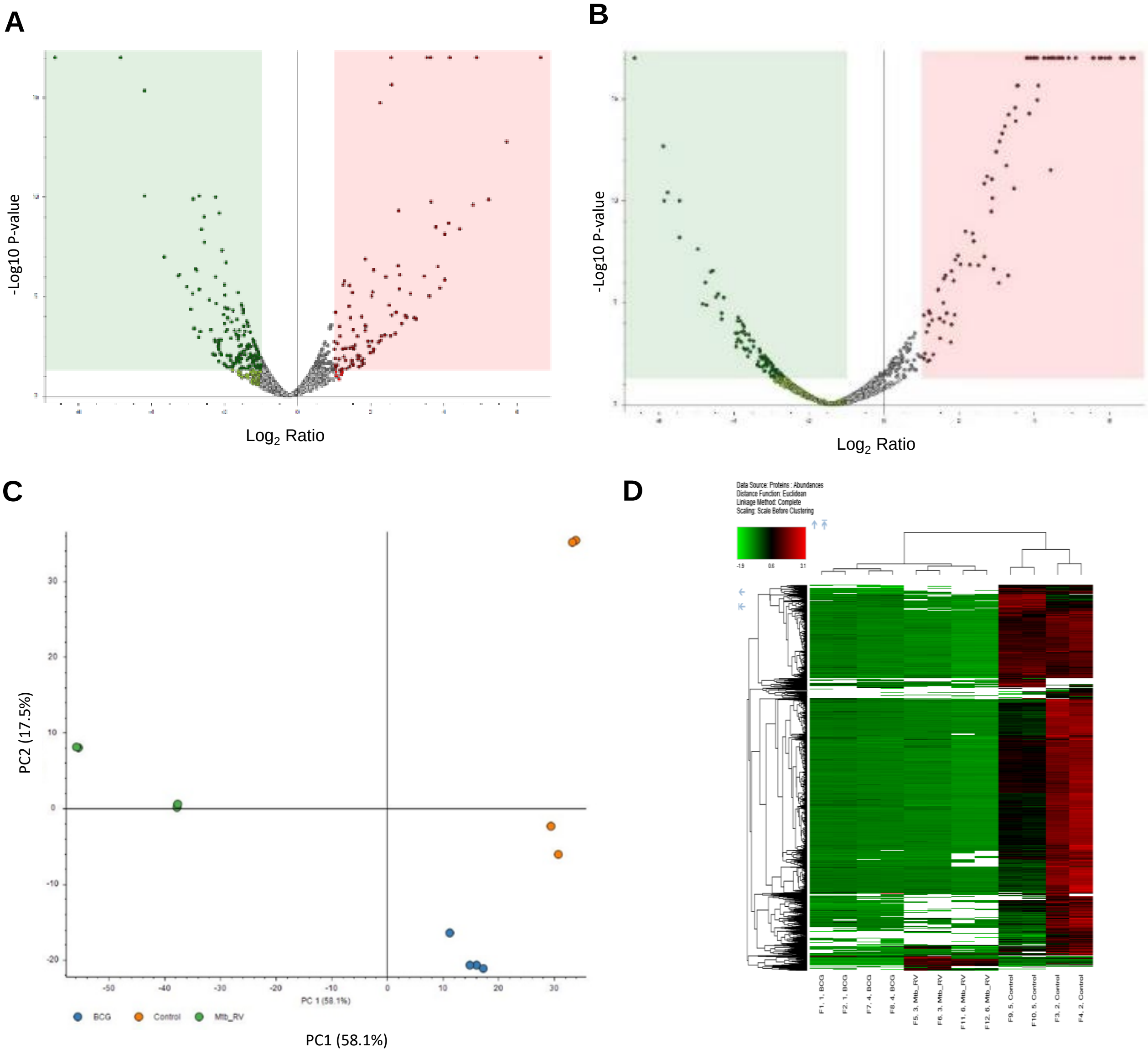


**Supplementary Figure S5. Comparative proteomic analysis of BCG and *M. tuberculosis* infected pulmospheres**.

A,B Volcano plot depicts log2fold change of up and down-regulated proteins in PS-BCG (A), and PS-*M.tb* in comparison with uninfected pulmospheres (B). Red and Green dots indicates up and-down regulated proteins, respectively; FDR cutoff is <0.05 and Log2FC>1.

C Principle component analysis (PCA) of uninfected, PS-BCG and PS-*M.tb* infected pulmospheres. PC1 and PC2 represents the percent of variance between (58.1%) and among (17.5%) the sample groups, respectively

D Hierarchical clustering analysis of peptide abundances in 3D vs 2D samples.

### Supplementary Figure S6

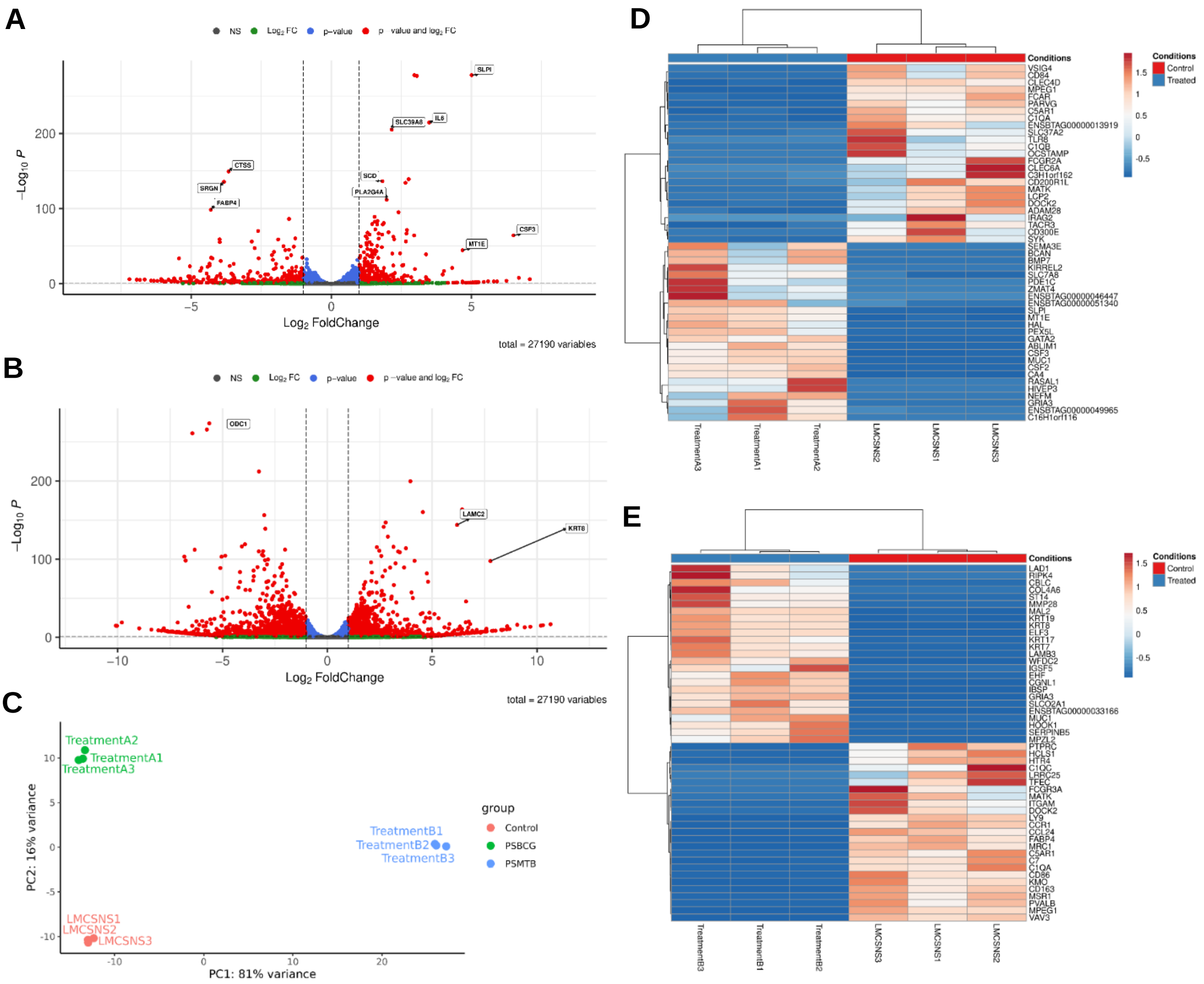


**Supplementary Figure S6. Comparative transcriptomic analysis of BCG and *M. tuberculosis-infected* pulmospheres**.

A,B Volcano plot depicts $log_2$fold change of up and down-regulated proteins in PS-BCG (A), and PS-*M.tb* in comparison with uninfected pulmospheres (B). FDR cutoff is <0.05 and $Log_2FC>1$.

C Principle component analysis (PCA) of uninfected, PS-BCG and PS-*M.tb* infected pulmosphere. PC1 and PC2 represents the percent of the variance between (81%) and among (16%) the sample groups, respectively.

D,E Heatmaps showing the top 50 up and downregulated DEGs in PS-BCG (D) and PS-*M.tb* groups (E).

### Supplementary Figure S7

#### A DEGs: BCG vs uninfected- biological pathways

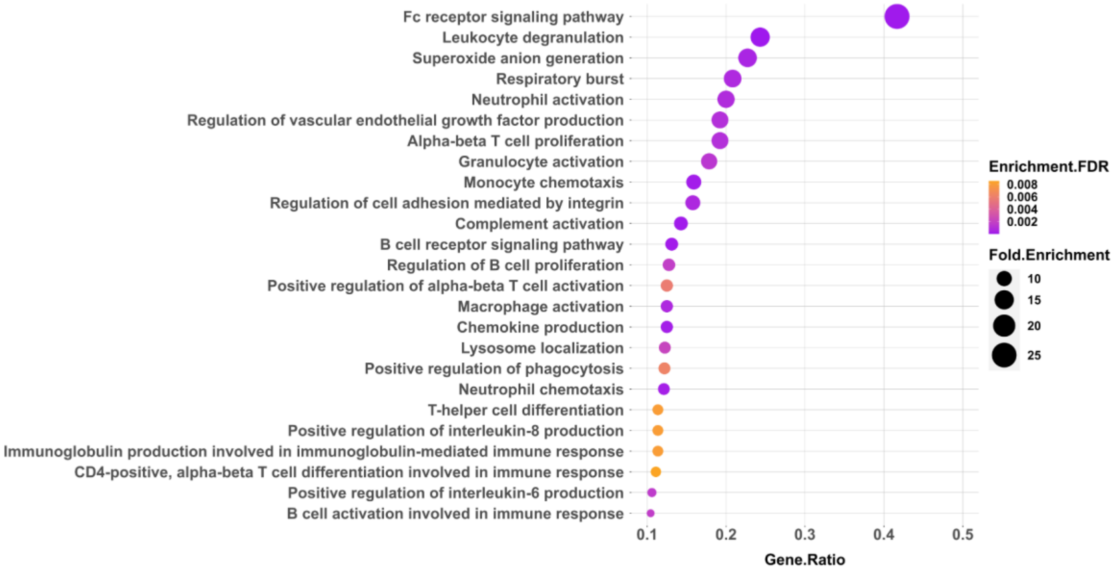


#### B DEGs: BCG vs uninfected- KEGG pathways

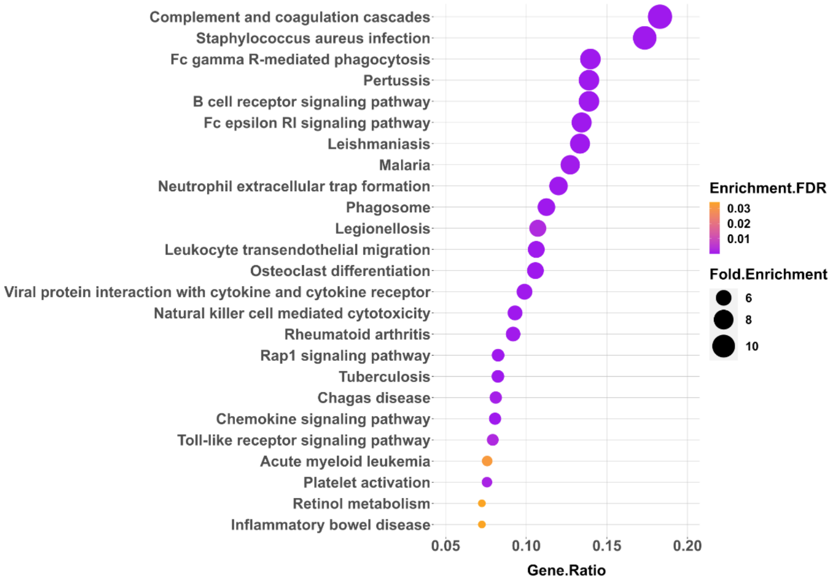


**Supplementary Figure S7. Functional enrichment analysis of downregulated DEGs in case of BCG vs uninfected pulmospheres**.

A, B Biological process (A), KEGG pathway analysis (B) of DEGs in PS-BCG. The X-axis represents the gene ratio of DEGs for each term relative to the total number of genes in that term, and the Y-axis represents the pathway. Dot size indicates the fold enrichment and emphasizes the degree to which the gene is strongly associated with the pathway. Dot color reflects the enrichment, as measured by the False Discovery Rate (FDR).

### Supplementary Figure S8

#### A DEPs: BCG vs uninfected- biological pathways

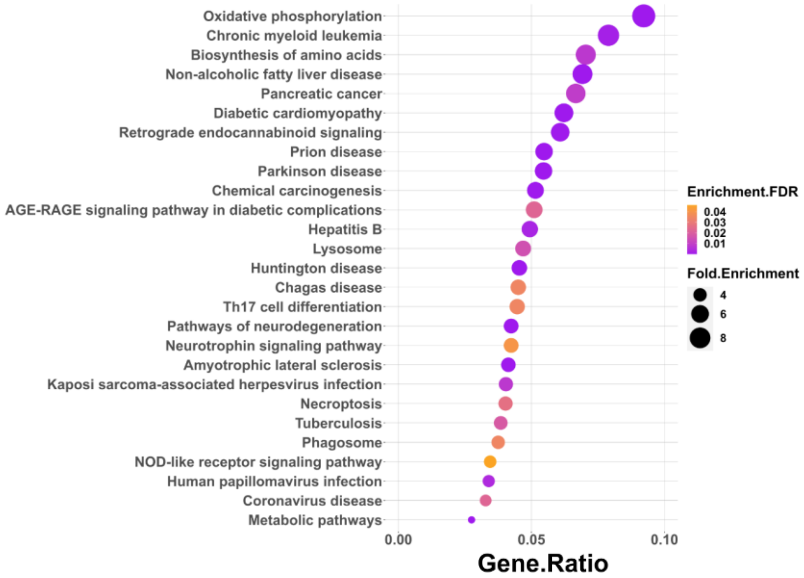


#### B DEPs: BCG vs uninfected- KEGG pathways

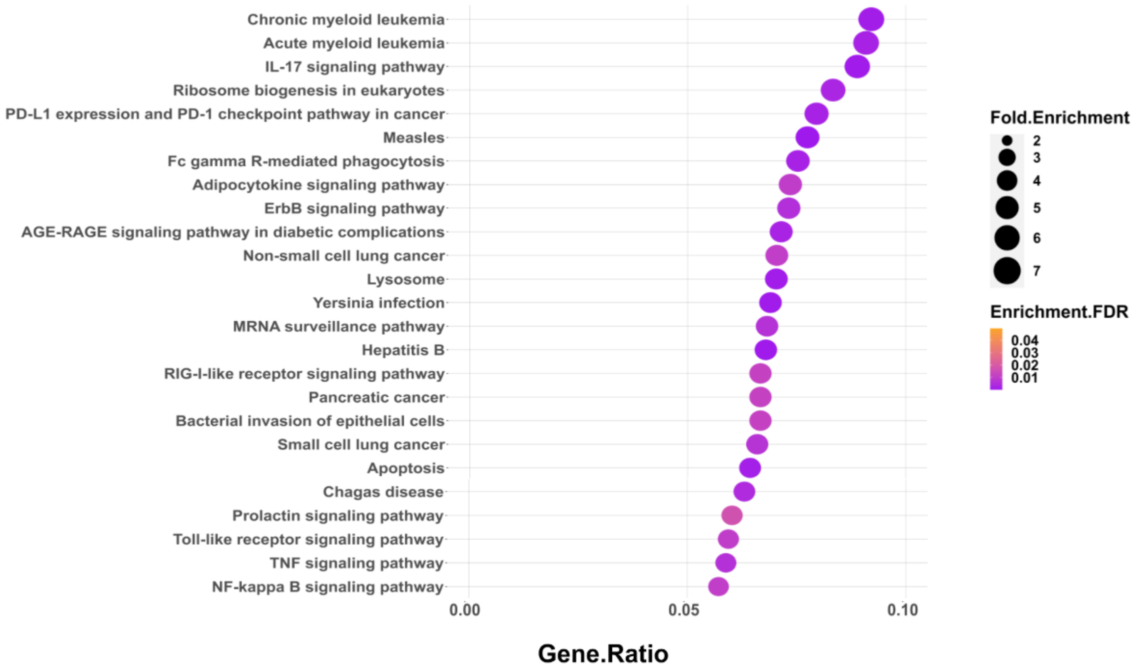


**Supplementary Figure S8. Functional enrichment analysis of down-regulated DEPs in case of BCG vs uninfected pulmospheres**.

A,B Biological process (A), and KEGG pathway analysis (B) of DEPs in PS-BCG. X-axis represent the gene ratio of DEPs for each term relative to the total number of genes in that term, and Y-axis represents the pathway. Dot size indicate the fold enrichment and emphasizes the degree to which the gene is strongly associated with the pathway. Dot colour reflects the enrichment, as measured by the False Discovery Rate (FDR).

### Supplementary Figure S9

**A** **DEGs: PS-*M.tb* vs uninfected- biological pathways**

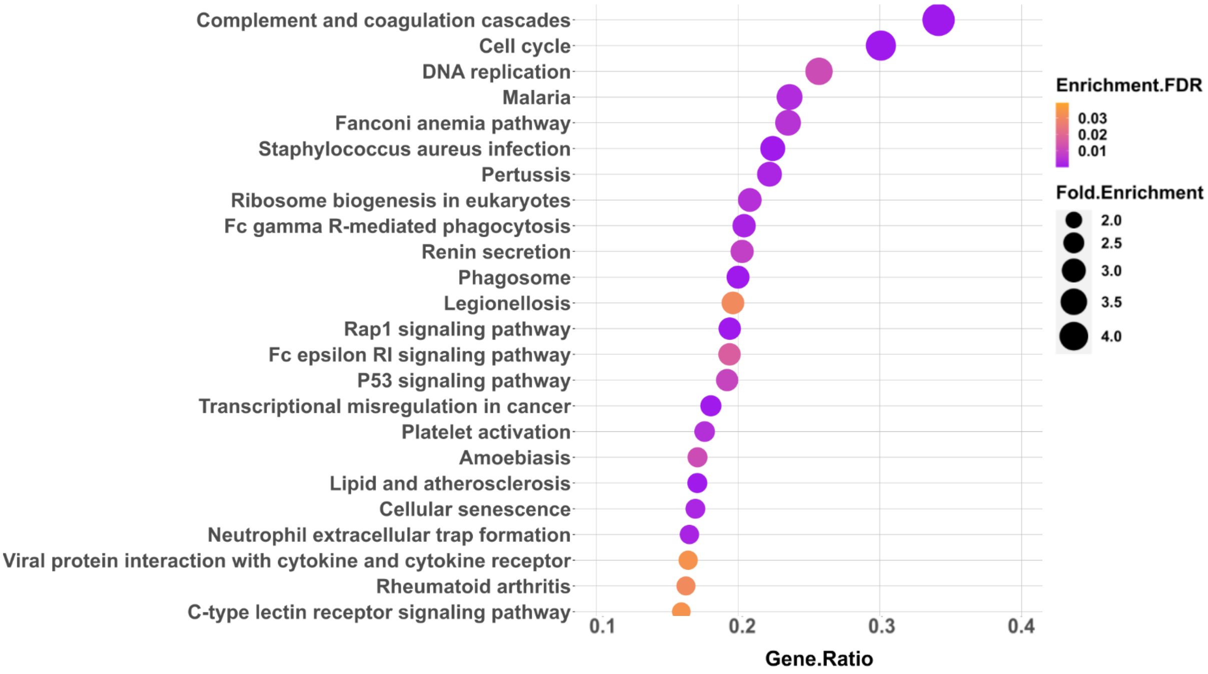


**B** **DEGs: PS-*M.tb* vs uninfected- KEGG pathways**

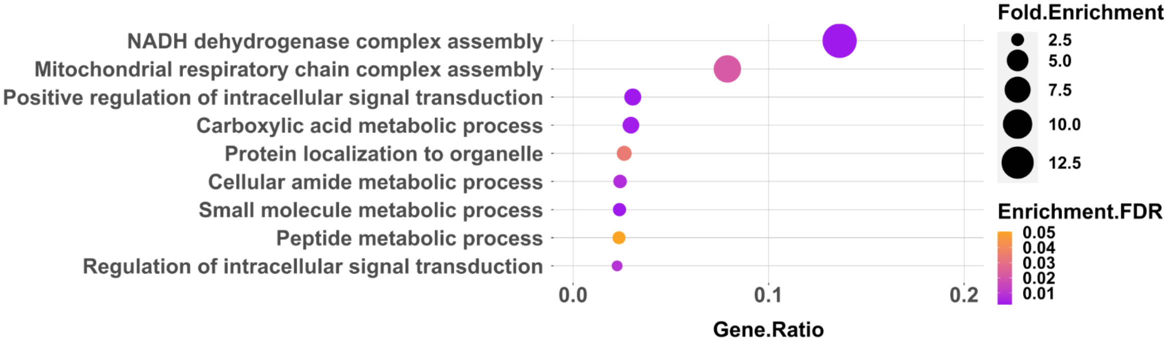


**Supplementary Figure S9. Functional enrichment analysis of downregulated DEGs in case of *M. tuberculosis*-infected vs uninfected pulmospheres**.

A, B Biological process (A), and KEGG pathway analysis (B) of DEGs in PS-*M.tb*. The X-axis represents the gene ratio of DEGs for each term relative to the total number of genes in that term, and the Y-axis represents the pathway. Dot size indicates the fold enrichment and emphasizes the degree to which the gene is strongly associated with the pathway. Dot color reflects the enrichment, as measured by the False Discovery Rate (FDR).

### Supplementary Figure S10

**A** **DEPs: PS-*M.tb* vs uninfected- biological pathways**

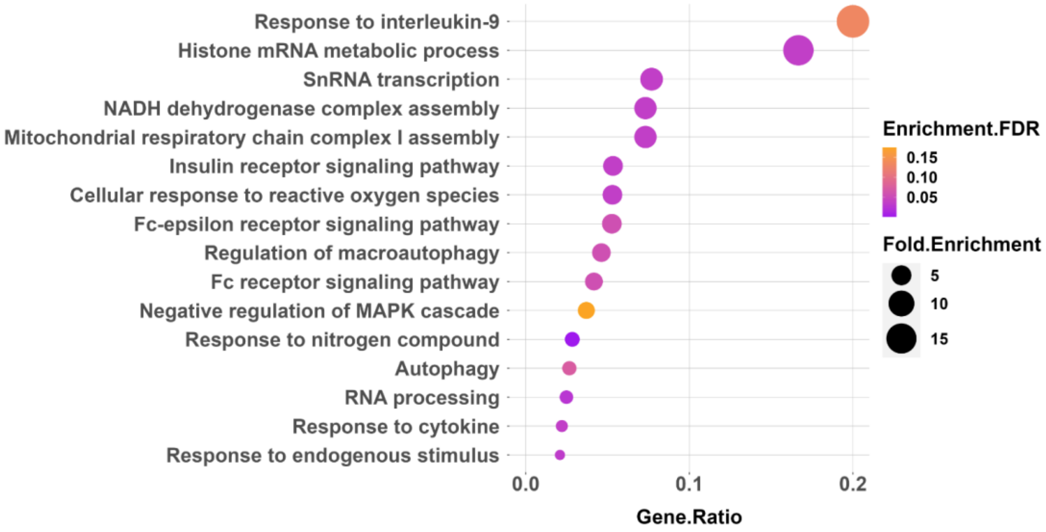


**B** **DEPs: PS-*M.tb* vs uninfected- KEGG pathways**

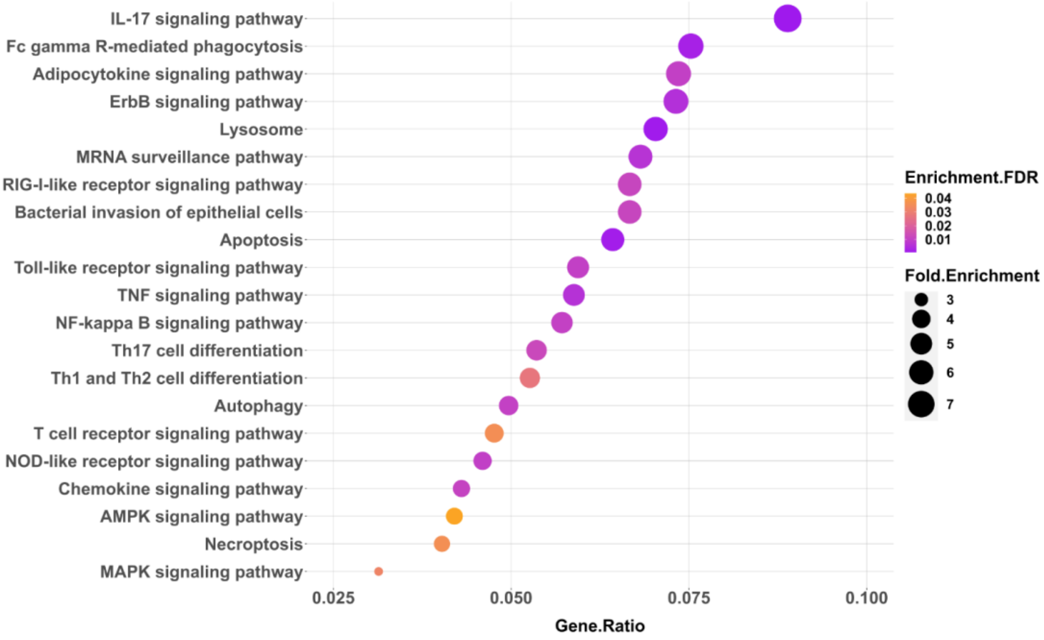


**Supplementary Figure S10. Functional enrichment analysis of downregulated DEPs in case of *M. tuberculosis*-infected vs uninfected pulmospheres**.

A, B Biological process (A), and KEGG pathway analysis (B) of DEPs in PS-*M.tb*. The X-axis represents the gene ratio of DEGs for each term relative to the total number of genes in that term, and the Y-axis represents the pathway. Dot size indicates the fold enrichment and emphasizes the degree to which the gene is strongly associated with the pathway. Dot color reflects the enrichment, as measured by the False Discovery Rate (FDR).
